# From Library to Landscape: Integrative Annotation Workflows for Compound Libraries in Drug Repurposing

**DOI:** 10.1101/2025.07.23.666341

**Authors:** Jeanette Reinshagen, Brinton Seashore-Ludlow, Yojana Gadiya, Anna-Lena Gustavsson, Ziaurrehman Tanoli, Tero Aittokallio, Johanna Huchting, Annika Jenmalm-Jensen, Philip Gribbon, Andrea Zaliani, Flavio Ballante

## Abstract

In the rapidly advancing landscape of drug discovery and repurposing, efficient access and integration of chemical and bioactivity data from public repositories has become essential. We implemented two complementary annotation pipelines (KNIME- and Python-based) designed to automate the extraction and integration of curated chemical and bioactivity data from public repositories. These pipelines are adaptable to any user-provided compound library, allowing reproducible workflows to integrate data from heterogeneous sources (e.g., ChEMBL and PubChem). As part of the REMEDi4ALL project, which aims to establish a European platform for drug repurposing, we validated our framework on a harmonized subset of the Specs repurposing collection (over 5000 compounds, available in-house). Additionally, we developed two interactive dashboards that support multilayered analyses and visualization by integrating chemical properties, bioactivity profiles, and relational data. We show how this framework streamlines the collection of harmonized data and facilitates analyses that are critical in drug repurposing efforts, while remaining versatile for broader applications in drug discovery. Both pipeline protocols are publicly available online, and the dashboards are open access.

## Introduction

Drug repurposing has emerged as a powerful strategy in drug discovery, offering a cost-effective and resource-efficient approach to identifying new therapeutic uses for existing drugs^1^. This method has become increasingly popular, especially for addressing unmet medical needs, e.g. in case of rare or orphan diseases and urgent health crises. Compared to traditional drug discovery, drug repurposing can reduce the development time from 10-17 to 3-12 years and costs from $2-3 billion to around $300 million per approved drug, while increasing the likelihood of success by up to 30%^2,3^. A notable example of successful drug repurposing is azidothymidine (AZT), which transitioned from an abandoned anti-cancer candidate to become the first FDA-approved drug for treating HIV infection^4^. Similarly, thalidomide which was once withdrawn from the market due to its teratogenic effects, was repurposed first for leprosy^5^ and later also for multiple myeloma^6^. Among more recent examples, lonafarnib, which was originally developed for the treatment of cancer, was approved as the first treatment for Hutchinson-Gilford progeria syndrome (HGPS)^7^.

Despite its advantages, data-driven drug repurposing still faces significant challenges. These include difficulties in accessing, integrating, and processing large volumes of heterogeneous chemical, physical and biological experimental data describing pre-clinical molecules, clinical candidates, and marketed drugs^8,9^. Critically, the success of these drug repurposing efforts relies on extensive biological and clinical data found in public repositories^10^, underscoring the need for standardization strategies and analytical methods. Currently, there are more than one hundred public and open databases in the biomedical domain, each covering distinct subjects such as genes, compounds, and diseases^11^. These resources serve as invaluable repositories of scientific knowledge, playing a pivotal role in advancing research and drug discovery. However, maintaining and updating these databases to keep pace with rapid scientific advancements requires dedicated teams of experts.

Repositories such as the Drug Repurposing Hub^12^, ChEMBL^13^, PubChem^14^, DrugBank^15^, Probes & Drugs^16^ and the Guide to PHARMACOLOGY^17^, among others, provide extensive information linking chemical structures, biological activities, mechanism of action and clinical data^18,19^. Since these databases continuously grow over time, they offer researchers an ever-expanding landscape of molecular and clinical information, allowing increasingly sophisticated drug repurposing strategies. However, the amount and complexity of available data requires modern computational approaches to effectively mine and interpret this information^20^. Indeed, advanced techniques in artificial intelligence and machine learning, along with relational databases, are increasingly being employed to uncover hidden patterns and relationships within these datasets^21,22^, potentially revealing novel drug-disease associations and accelerating the repurposing process^23,24^.

Despite their individual significance, these databases often remain siloed, limiting their potential to provide holistic insights into complex biological systems and diseases. To overcome this fragmentation, integrated workflows that combine data from multiple sources are essential. Such workflows can uncover hidden relationships among genes, compounds, targets, and diseases, enabling a more comprehensive understanding of biological mechanisms and facilitating translational research. However, significant challenges lie in establishing standardized protocols for data collection, curation, and sharing across the scientific community. A critical factor in the success of these integrative approaches is the selection of resources that adhere to the FAIR (Findable, Accessible, Interoperable, and Reusable) principles^25,26^. Databases compliant with FAIR principles ensure that data can be easily located, accessed, and efficiently integrated across platforms without loss of meaning or context. This compliance is crucial for efficient harmonization of datasets, reducing redundancy, and promoting data interoperability. In the era of data-driven research, the ability to efficiently harmonize and utilize vast biomedical datasets will be a key driver of innovation.

Harmonized datasets are indeed essential in modern scientific research, particularly in biomedical fields. They enable the integration of diverse data sources into a unified and standardized framework, when the FAIR principles are consistently applied. This standardization enhances the ability to validate citations, improves the statistical robustness of analyses, and allows researchers to evaluate the generalizability of findings across different contexts^27^. In drug repurposing, data harmonization significantly accelerates research timelines by streamlining data integration and analysis. Furthermore, harmonized datasets promote interoperability among diverse data sources, facilitating collaboration between researchers and encouraging knowledge sharing throughout the scientific community. For instance, the Alzheimer’s Disease Neuroimaging Initiative (ADNI) has contributed to over 600 publications on Alzheimer’s biomarkers, underscoring the value of standardized datasets^28^. In healthcare, harmonized clinical data enables more precise analyses and diagnoses, facilitates personalized treatments, and enhances the efficiency of AI models^29^. Despite these benefits, challenges such as data heterogeneity, ethical concerns, technical barriers, and varied regional regulations persist^30^. Overcoming these obstacles requires advanced frameworks and universal standards, such as HL7 Fast Healthcare Interoperability Resources FHIR^31^, an interoperability standard enabling heath data exchange between different software systems, to fully harness the potential of harmonized data in driving scientific discovery and innovation.

Despite the development of various methods for collecting and analyzing annotated data from screening libraries^32–36^ the exponential growth in data volume and complexity requires continued innovation in this field. Accordingly, the development of up-to-date methods remains critical in drug repurposing. Moreover, there is a need for an easy-to-use standardized method that can be widely adopted across different research settings. Such a protocol would democratize access to powerful data processing and analysis techniques, enabling researchers from diverse backgrounds to effectively explore the information available from chemical repositories.

In this work, we present a framework aimed at advancing drug repurposing efforts through the development of pipelines for annotating compound screening libraries, as well as platforms for visualization and multilayered analysis of annotated data. The developed workflows are explicitly designed to facilitate automated integration and interpretation of underlying data using dynamic approaches, ensuring alignment with the latest available information. We demonstrate the applicability and reusability of this framework using the Specs Repurposing Library^37^, which is a subset of commercially available compounds described in the Broad Institute’s Drug Repurposing Hub^38^. This approach provides solutions for both computational researchers and non-computational scientists, offering robust and practical tools to leverage prior knowledge effectively for their drug repurposing projects. By adhering to FAIR principles, the method ensures high-quality curated data, as well as easy access and reusability. Although the present research focuses on molecules with clinical trial history, aiming to provide a useful resource for informed decision-making in drug repurposing, the reported pipelines can be readily adapted to different drug discovery projects.

Aimed at serving a broad community, we developed two distinct complementary platforms: one based on Python and NEO4J^39^ and another utilizing KNIME Analytics Platform^40^ (See Data Availability Section).

This work is part of a collaborative initiative within the REMEDI4ALL EU project^41^ involving scientists from the Fraunhofer Institute for Translational Medicine and Pharmacology (ITMP) and the Karolinska Institute (KI) who have collaborated to unify their compound collections under a shared identification procedure.

## Results

We designed and developed two distinct pipelines to gather, process, and analyze annotated data from publicly available repositories. Among others, the collected data included key annotations, such as compound identifiers, structural information, physicochemical properties, bioactivity profiles, and metadata related to biological targets, assay conditions, therapeutic use, research and patent literature. We applied both approaches to the Specs repurposing set, a well-established collection of compounds with potential for drug repurposing. This served as a test case to showcase the pipelines’ capabilities in collecting, processing, structuring, and analyzing bioactivity data in a drug repurposing context. Notably, the two pipelines can be applied to any chemical library provided by users, making them versatile tools for various drug discovery projects. To facilitate interaction with the collected results, we developed two complementary analytical dashboards. The first offers real-time annotation and analysis capabilities for any single chemical compound input, while the second focuses specifically on the Specs repurposing set, allowing efficient navigation and analysis of complex relationships in a graph structure.

### Development of KNIME and Python pipelines

A KNIME (Konstanz Information Miner) pipeline, named “R4A Annotation Tool”, was developed as a graphical workflow for data extraction and processing. This workflow to annotate small molecules with chemical, bioactivity and drug information takes the compound InChIKey as input (Figures 1a and 16). The InChIKey is a highly detailed and unique representation of a chemical structure, making it an ideal input for our process. The next step involves mapping the InChIKey to chemical identifiers using PubChem. This step allows us to gather all known identifiers for the compound, including the Compound Identifier (CID), ChEMBL Identifier (ChEMBL ID), and SureChEMBL Identifier (sChEMBL ID). Using the ChEMBL IDs, we query the ChEMBL database to extract four key levels of information. The first level includes physicochemical properties, such as molecular weight, preferred compound name, and structural features like hydrogen bond donors and acceptors, LogP, and etc. The second level focuses on bioactivity annotations. We prioritize binding and functional assays against single protein targets and retrieve descriptions and results for experiments, filtered by a confidence score of 8 or higher in ChEMBL. Additionally, to the target information (Target ID, Target pref name), we retrieve potency indicators such as IC_50_, EC_50_, Ki, Kd, CC_50_, and LD_50_, summarized by the pChEMBL value, which is the negative logarithm of the concentration value. This semi-positive defined value allows for easy numerical ranking of compound activity, with higher values indicating greater potency^36^. The third level provides drug approval metadata, including the year of first approval, maximum clinical phase achieved, and any withdrawal status. Finally, the fourth level includes clinical trial annotations along with their respective drug indications. Consistent ontologies such as the Experimental Factor Ontology (EFO) and Medical Subject Headings (MeSH) are used to represent indication areas. Additionally, we gather information on whether the compound has progressed through clinical phases and, if so, the latest clinical stage reached. The workflow described above is re-used for the creation of an interactive dashboard version that handles single compound input. Here, the compound’s ChEMBL ID is used as input and future versions will expand to also allow querying with machine-readable chemical structure representations (SMILES, InChIKey) or trivial names as input. The application gives users the possibility to set constraints for confidence score range, minimum pChEMBL value and assay type (binding and/or functional). For institutions with access to KNIME Business Hub, the dashboard is available as web application to be used without KNIME installation.

**Figure 1.**
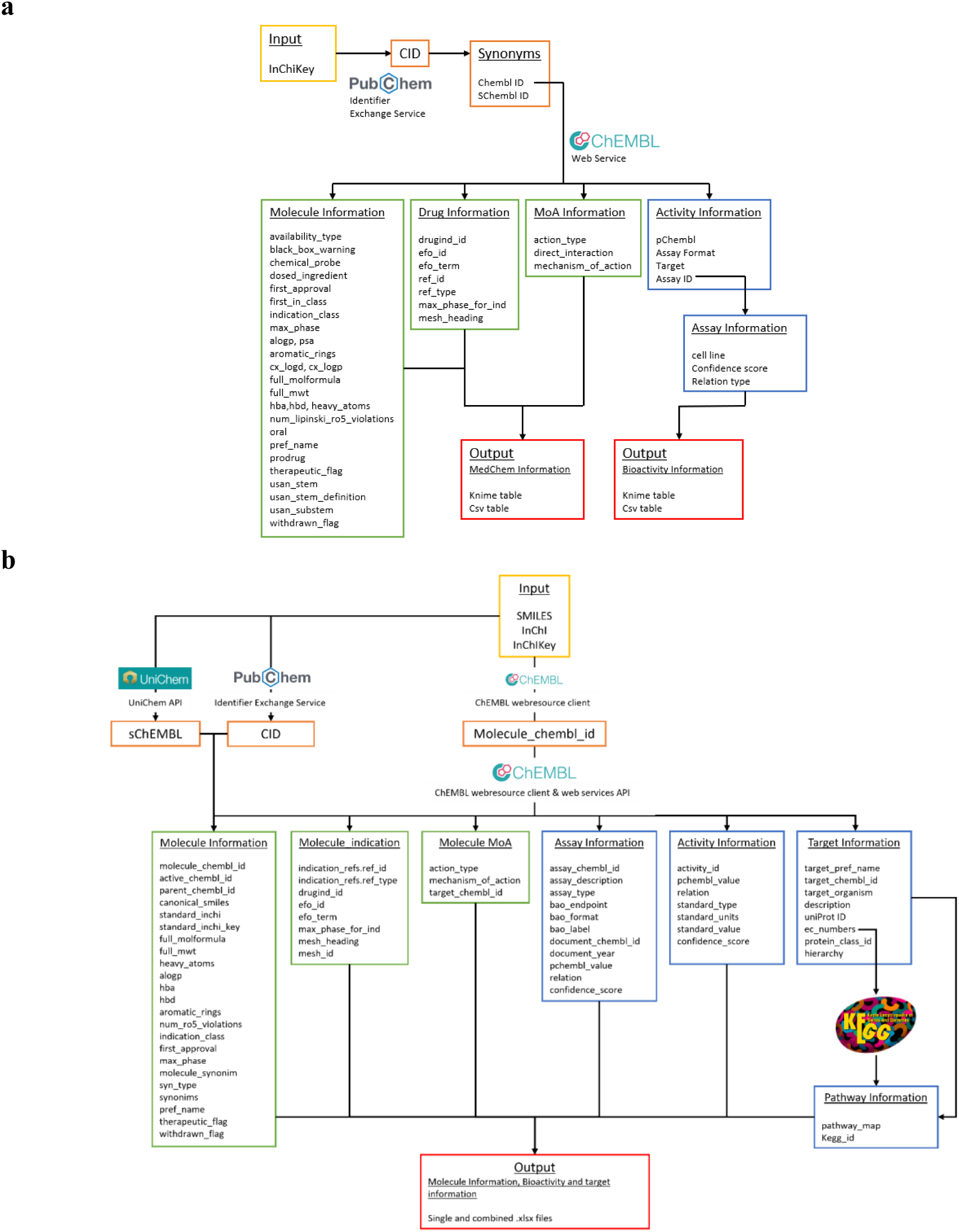
Overview of the KNIME and Python annotation workflows. a) KNIME pipeline: in the workflow, we make use of both PubChem and ChEMBL databases to aggregate all relevant biological and chemical information, b) Python pipeline: different molecular identifiers (SMILES, InChI, InChIKey) can be used as input to query the ChEMBL, PubChem and UniChem databases and extract relative compound identifiers. These identifiers allow access to relevant molecular, assay, activity, pharmacological, target and pathway information, which are compiled in output .xlsx files.

The KNIME pipelines offer full integration capability into user-specific drug discovery analyses, both downstream of hit identification workflows, e.g. or upstream of machine learning or SAR studies. The KNIME workflow offers a wide variety of export and import formats to integrate with users’ infrastructure, e.g. MS Excel, SDF or Tibco Spotfire.

A Python pipeline, named “Chemical Annotator”, was implemented as a script-based solution (Figure 1b and Supplementary Figure S1), for systematic retrieval, integration, and organization of chemical, bioactivity, and target information. It accepts various chemical notations as input, including SMILES, InChI, or InChIKey, which are processed through three resources: ChEMBL, UniChem, and PubChem. These resources are used to obtain corresponding molecule identifiers such as ChEMBL ID, PubChem CID and sChEMBL. The workflow mainly relies on the molecule ChEMBL ID to extract relevant metadata from ChEMBL. This data includes chemical identifiers (such as SMILES, InChi, InChIKey and molecular formula), molecular properties (including molecular weight, number of heavy atoms, AlogP, hydrogen bond donors and acceptors, number of aromatic rings, and number of Lipinski’s rule of 5 violations), as well as synonyms, approval date, withdrawal status, and therapeutic classification. Clinical and pharmacological context is also included, such as drug indications, clinical phase and associated disease terms, for example, EFO terms and MeSH headings. The Chemical Annotator further collects data on mechanisms of action, target identifiers, experimental assay details and quantitative bioactivity measurements. Additionally, it gathers target-related information, including target names, ChEMBL and UniProt identifiers, organism, enzyme commission numbers and protein class hierarchies. Associations with biological pathways are retrieved from the KEGG database^42^. All collected data are organized and exported in both single and combined .xlsx files (Figure 1b).

Both pipelines and demonstration data outputs are publicly accessible (see data and code availability sections).

### Development of KNIME and Neo4j dashboards

To offer a comprehensive approach for visualization and analysis of the collected data from the two annotation pipelines, we built both KNIME and Neo4j dashboards. The KNIME dashboard uses the KNIME-based annotation workflow to annotate individual compounds on-the-fly by querying the relevant databases directly, designed to generate a quick overview of most relevant and up-to-date information, while the Neo4j dashboard specializes in graph-based visualizations tailored to the Specs repurposing set, ideal for exploring complex relationships in drug interactions and biological pathways. This dual approach allows researchers to gain insights from statistical analysis and network visualization, enhancing decision-making in drug repurposing and discovery studies.

### KNIME dashboard

The “ChEMBL Annotation Dashboard” is the KNIME-based dashboard version of the annotation workflow. Upon querying single substances, it displays various visualizations for chemical, physicochemical, pharmaceutical, clinical trial and bioactivity information retrieved directly from the ChEMBL database.

The user of this data app can set constraints for the target confidence range, minimum pChEMBL value, and assay type using the drop-down menus next to the ChEMBL ID input field. Figure 2 exemplifies the visualizations available using Diclofenac Sodium (CHEMBL1034) as input, displaying the compound’s structure, alongside physicochemical information, a time series on reported activity, frequency of reported targets as pie diagram and statistics on reported potency in a boxplot.

**Figure 2.**
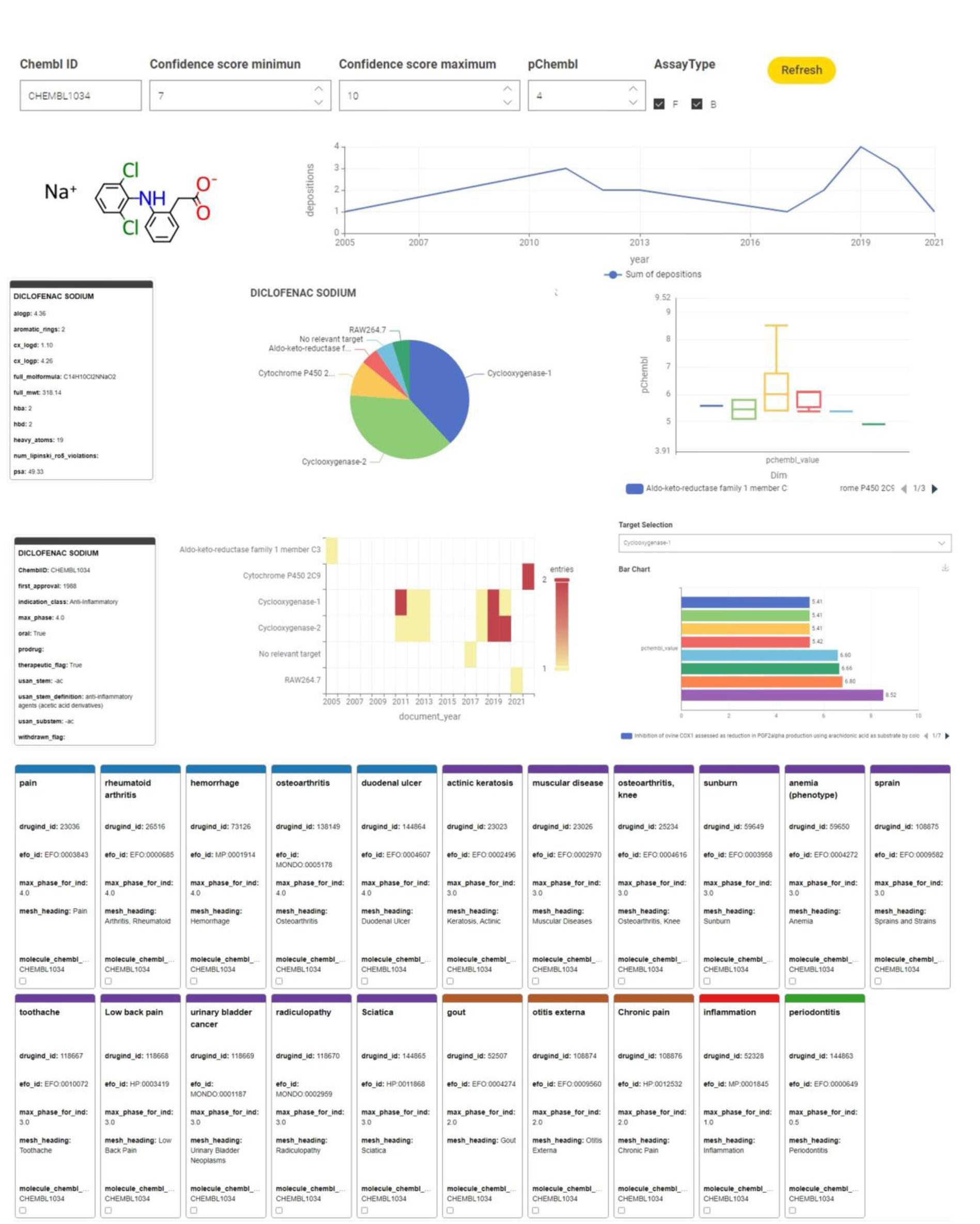
Example dashboard output using the KNIME annotation tool for a single compound, Diclofenac sodium.

Further visualizations offer a comprehensive overview of the pharmaceutical information, including the year of first approval, the therapeutic indication class, oral administration possibility and whether the drug was withdrawn. The frequency per year the compound was reported as being active (as defined by the user’s set constraints) against a specific target is illustrated in a heatmap and in more detail, the associated pChEMBL values for the specific user selectable targets is presented in an interactive bar chart, where a drop-down menu lets the use select the target of interest. The indication area, EFO ID and status of ongoing and completed clinical trials in ranked information tiles complete the dashboard.

### Neo4j-powered dashboard

We named the Neo4j powered dashboard “Chemical Biology Atlas”. The Chemical Biology Atlas provides a versatile and comprehensive platform for the exploration and analysis of chemical biology data. The dashboard is composed of multiple interactive panels, *Compound search*, *Compound’s analogues space search* and *Target search* (Figure 3). Each page contains a collection of self-contained visual or interactive elements (namely reports), which display specific information. In addition, displayed data can be downloaded as comma separated values (CSV) file.

**Figure 3.**
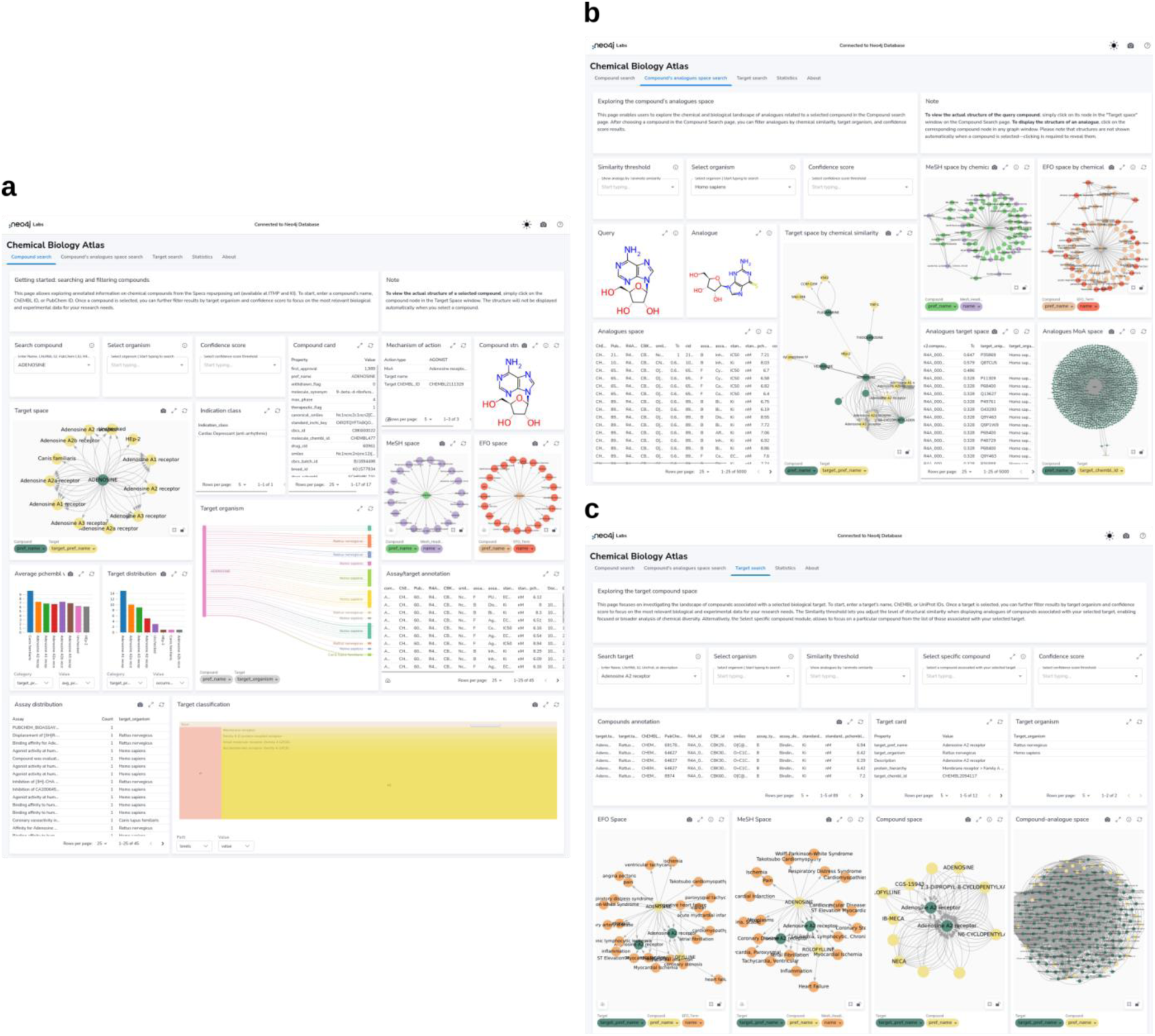
Neo4j Dashboard. a) Compound search panel; b) Compound’s analogues space search panel; c) Target search panel.

The *Compound search* page (Figure 3a) provides reports for exploring and analyzing compound data. Users can retrieve detailed information about compounds using various identifiers such as ChEMBL ID, PubChem CID, InChIKey, and more. The *Compound card* report provides a detailed overview of compound’s properties including identifiers, molecular structure, clinical or preclinical records. The *Mechanism of action* report details the biological mechanisms of the compound, including action types and targets. The *Compound structure* report depicts the compound structure. The *Target space* report provides insights into the interactions between compounds and their targets in a form of interactive network graph map. In a similar fashion, relationships between the compound of interest and biomedical ontologies (i.e., EFO and MeSH) can be explored through the *EFO space* and *MeSH space* reports, respectively. The *Target organism* report provides an interactive Sankey diagram visualizing relationships between compound, activities, targets and target organisms. The *Select organism* report allows users to select target organisms to perform focused searches and analyses on specific organism. The *Average pChEMBL value* report displays the average pChEMBL value for each target, offering a measure of half-maximal response of the compound of interest against a specific target. The *Target distribution* report shows the distribution of targets associated with the compound, helping in understanding their prevalence and diversity. The *Assay/target annotation* report offers detailed information about assays and targets. The *Assay distribution* report highlights the types and frequencies of reported assays. The *Target classification* report shows a structured view of target types based on protein hierarchy, and the *Indication class* report provides information on the therapeutic areas associated with the compound of interest. Users can also filter results based on the C*onfidence score* assigned to the bioactivity data.

The *Compound’s analogues space search* page (Figure 3b), allows users to explore analogues or dissimilar compounds, compared to the molecule of interest, to analyse their biological activities and relationships with targets. A key feature is the ability to fine-tune the selection of analogues by adjusting the Tanimoto similarity index through the *Select similarity threshold* report. The *Target space by chemical similarity* report enables the identification of such compounds and their associated target information, which can be filtered by target organism through the *Select organism* report, providing insights into potential shared biological activity and off-target effects. The *MeSH Space by chemical similarity* and the *EFO Space by chemical similarity* reports show MeSH heading and EFO terms based on chemical similarity, aiding in the classification and exploration of disease-related terms, respectively. The *Analogues MoA space* report offers detailed information about the biological mechanisms of compounds with certain similarity from the compound of interest. The *Analogues space* and *Analogues target space* reports provide insights into biological activity and targets of compounds ranked by their chemical similarity to the compound of interest. Users can also filter results based on the C*onfidence score* assigned to the bioactivity data.

The *Target search* page (Figure 3c) allows users to retrieve detailed information about targets using various identifiers such as ChEMBL ID, UniProt ID, and descriptions. The *Target card* and the *Target organism* reports provide comprehensive information and overview of the organisms associated with the selected target, respectively. The *Compound space* report displays compounds associated with the target of interest, while the *Compound-analogue space report* allows to explore analogues, and their targets, of the compounds associated with the selected target. The *MeSH space* and the *EFO space* reports display MeSH heading and EFO terms, respectively, related to the compounds associated with the target of interest and their analogues. The *Select similarity threshold* report allows to define a Tanimoto index value for analogue selection. Results can be further refined by applying the *Select organism* and *Confidence score* filters, in the same way as previously described. The *Select specific compound* report allows to focus on a particular molecule from the list of those associated with the selected target.

The *Statistics* page provides a measure of the dataset size and diversity, including the count of distinct compounds, targets, ligand-target activities, chemical similarity relationships, sureChEMBL IDs, ChEMBL IDs for compounds and target, UniProt entries, indication classes, mechanism of action and PubChem CIDs.

### Annotating the Specs repurposing set using both pipelines

We utilized the Specs repurposing library, which is physically housed at both ITMP and KI institutes. The first step in our efforts involved harmonizing the compound IDs between the two institutions’ collections to ensure consistency when integrating data from both sites and facilitate collaboration. This process culminated in a curated library representing the overlap between ITMP and KI collections and in the generation of a unique Remedi4All identifier (R4A ID) for each compound. Harmonization provided a common “Remedi4All harmonized repurposing set (R4A set) of 5254 entries (5230 unique substances), representing 93% of the original Specs^37^ set, as primary input for the annotation pipelines, with unified reference point for subsequent analyses (see Methods section). The KNIME and Python protocols were both used to extract and process data for the R4A set by using chemical structures as query. We focused our analysis on assay types B and F (binding and functional), with confidence scores ≥ 8 and pChEMBL values ≥ 6, focusing on most active substances with activity reported on defined targets. Thereby, we are filtering the available data to results of well-defined experimental conditions, where direct interaction with a specific and clearly identified target of interest is determined. Adjustment of these constraints is recommended depending on the specific scientific question and can be easily customized by the user in both pipelines. Despite employing distinct workflows for compound annotation (Figure 1), the two pipelines demonstrate comparable effectiveness in processing large datasets and notable alignment in their results, showing closely comparable outcomes across diverse metrics (Figures 4-8 and Discussion section). Importantly, both approaches keep track of the data sources together with registry IDs, such as ChEMBL IDs that have been assigned to compounds, assays, and targets. This enables users to trace back the original data source at any point to confirm data quality and further support FAIR data re-use.

**Figure 4.**
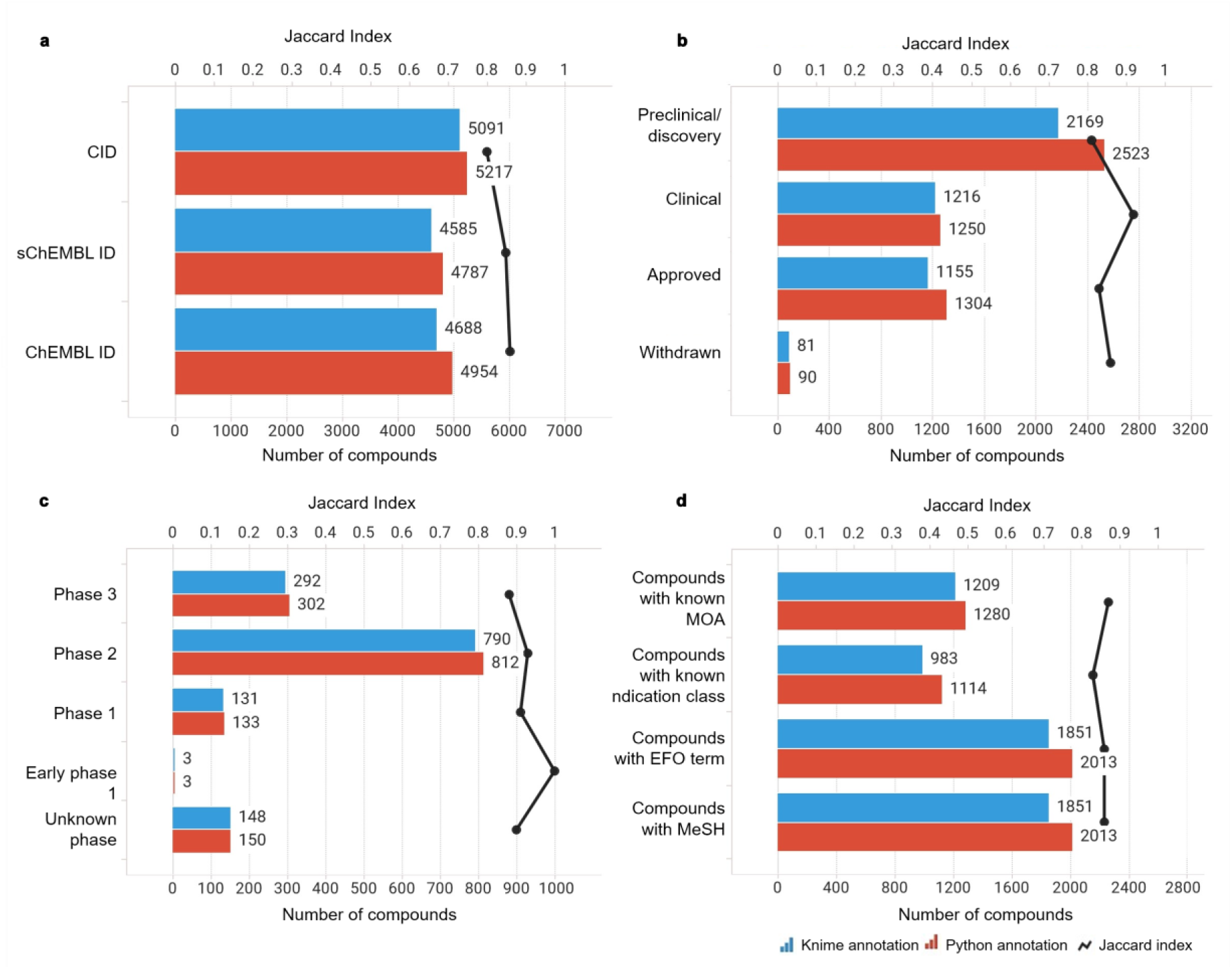
Compound-centric information retrieved by the KNIME and Python annotation pipelines (blue and red bars, respectively). The similarity between the two sets was measured using the Jaccard Index (black line). Data from binding and functional assays with a pChEMBL value ≥ 6 and a confidence score ≥ 8 were considered.

**Figure 5.**
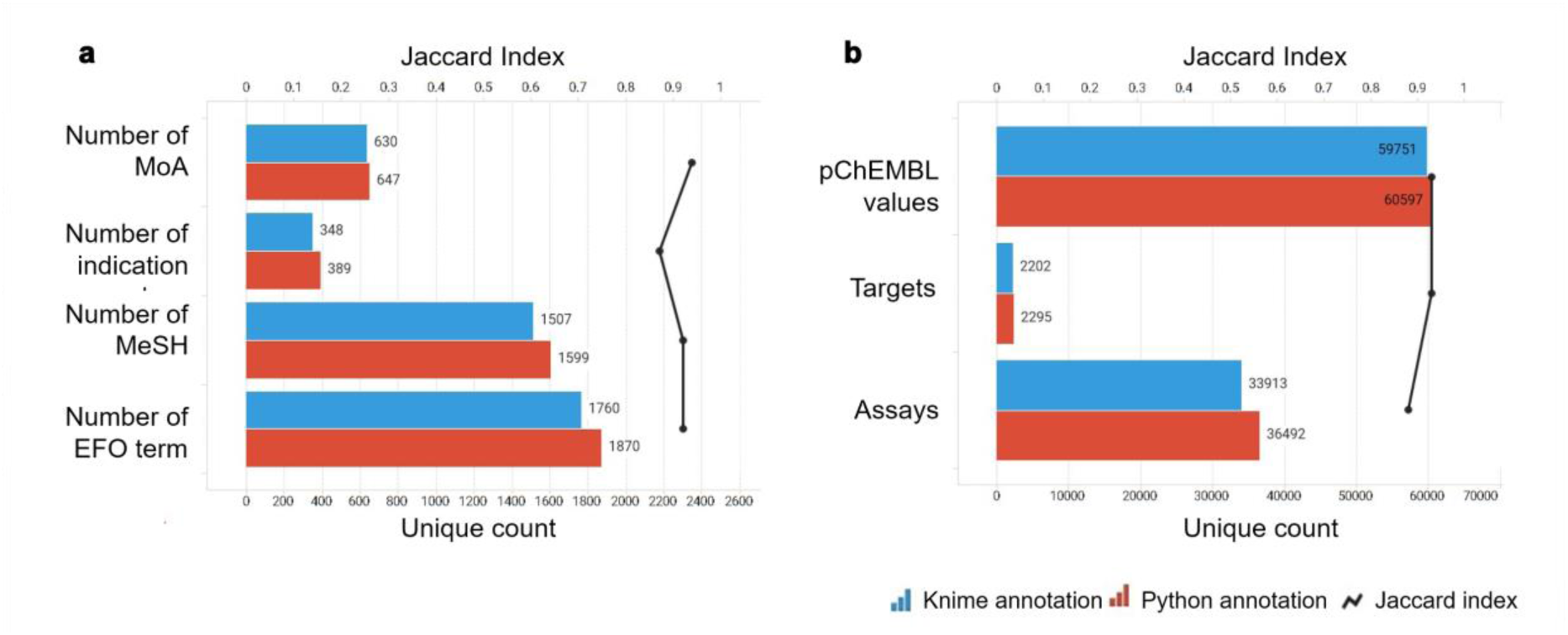
Annotation-centric information retrieved by the KNIME and Python annotation pipelines (blue and red bars, respectively). The similarity between the two sets was measured using the Jaccard Index (black line). Data from binding and functional assays with a pChEMBL value ≥ 6 and a confidence score ≥ 8 were considered.

**Figure 6.**
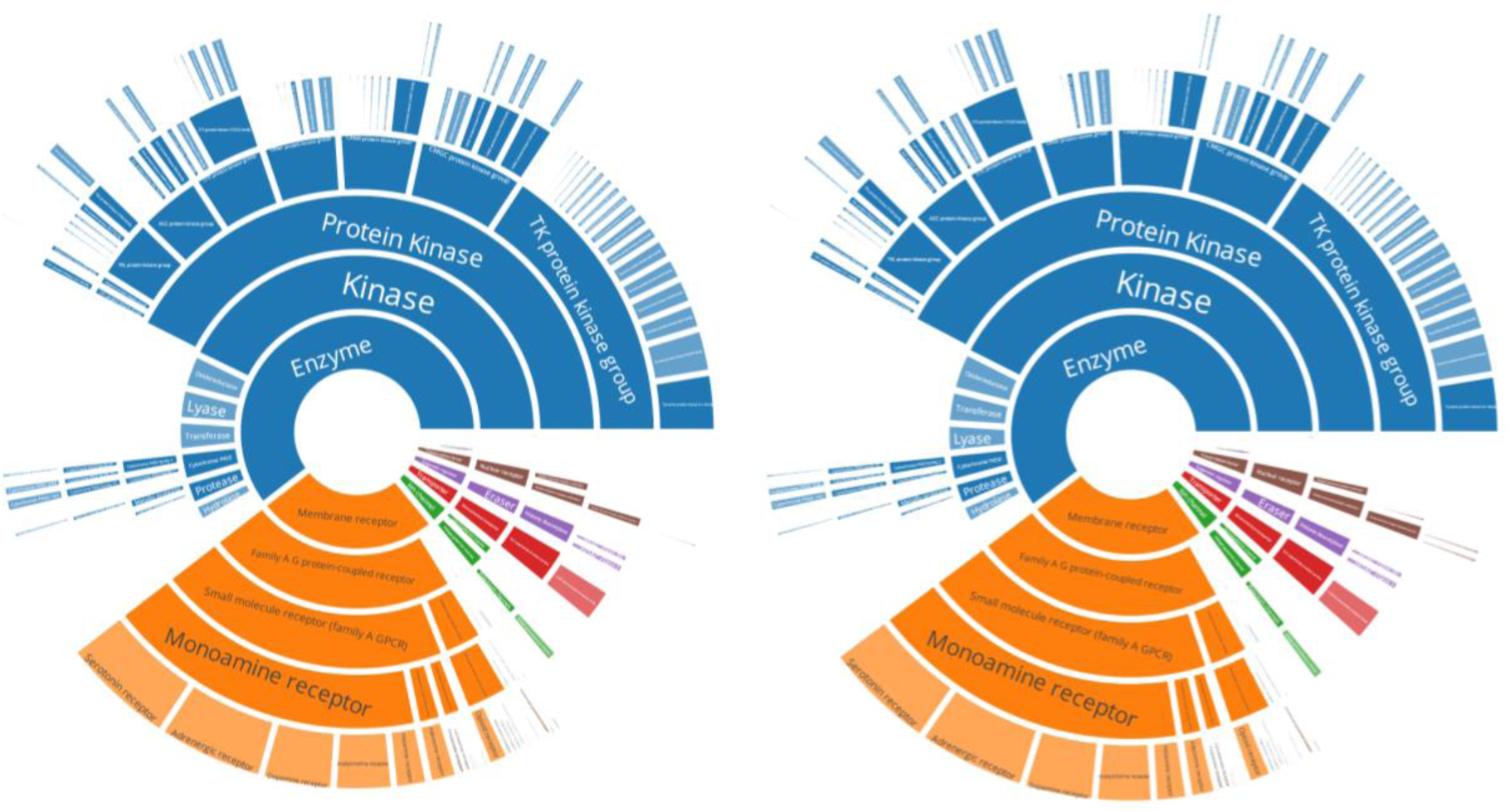
a) ChEMBL protein family classification from KNIME’s annotation; b) ChEMBL protein family classification from Python’s annotation. Data from binding and functional assays with a pChEMBL value ≥ 6 and a confidence score ≥ 8 were considered.

**Figure 7.**
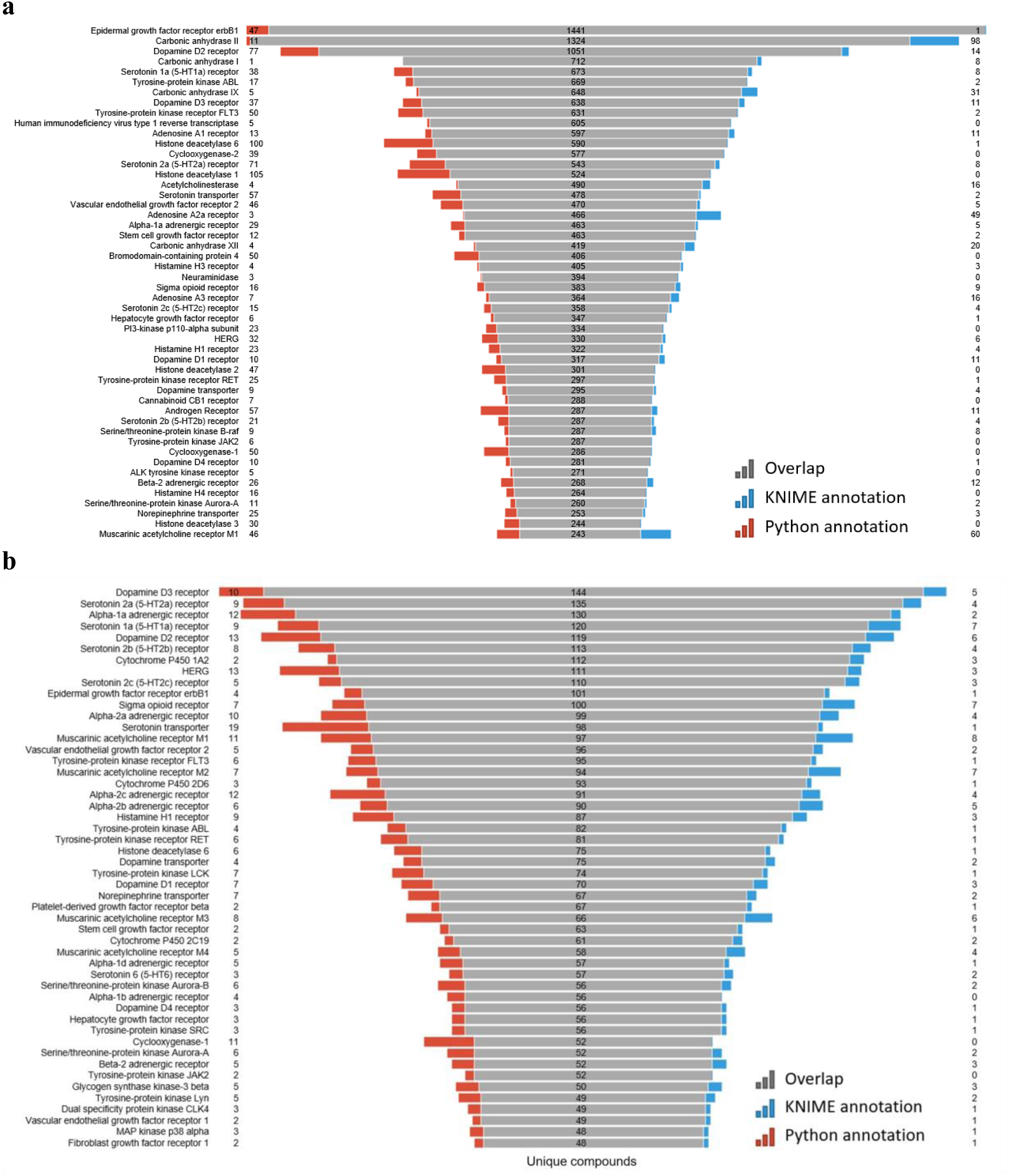
a) Most frequently active reported targets in ChEMBL database for the R4A set (i.e., number of pChEMBL values per target); b) Top-ranked targets based on n of unique compounds. Data from binding and functional assays with a pChEMBL value ≥ 6 and a confidence score ≥ 8 were considered.

**Figure 8.**
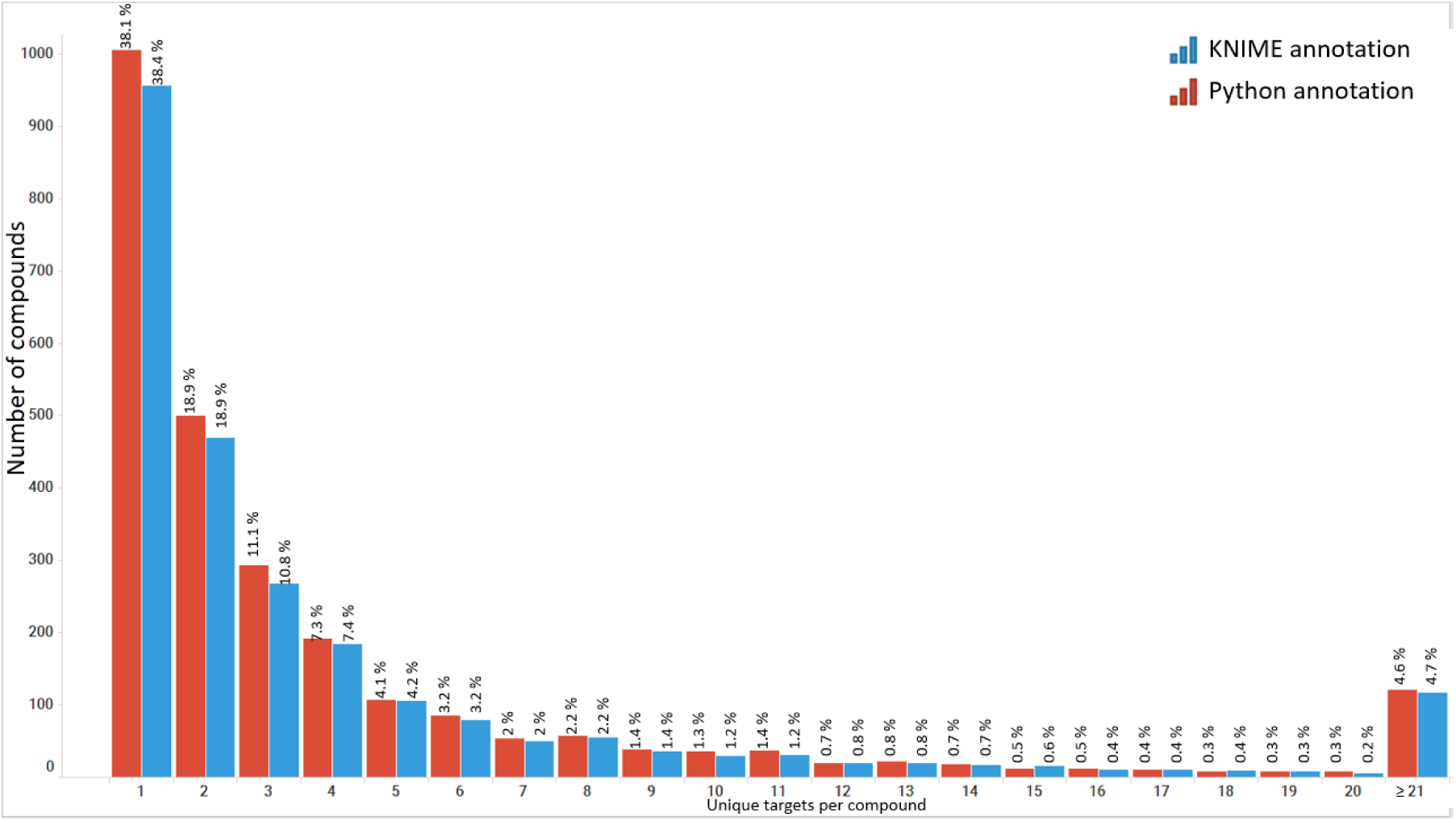
Distribution of compounds from the R4A set indicating the number of targets in Homo sapiens they are reported to be active against. Data from binding and functional assays with a pChEMBL value ≥ 6 and a confidence score ≥ 8 were considered.

### Applicability of a harmonized dataset for improved drug repurposing

We tested the applicability, strengths and weaknesses of the pipelines and dashboards with three use cases relevant to drug repurposing. The first use case is focused on a specific chemical substance, Tazarotene, enabling us to gather comprehensive data and insights tailored to this pharmaceutical uncovering its potential and hurdles for reuse in other indication areas. The second use case involved exploring the ion channel target KCNQ2 and the elucidation of the mechanism of action of a potent agonist found in a high-throughput screening campaign using the R4A set. The third use case demonstrates the use the Chemical Biology Atlas for inferring off-target effects of small molecules based on the annotations of close structural analogues.

#### Use case 1

Multiple Sulfatase Deficiency (MSD) is an ultra-rare lysosomal storage disorder with birth or juvenile onset. It is caused by mutations of the SMUF1 gene encoding the formylglycine-generating enzyme (FGE) leading to reduction of cellular sulfatase activities. MSD is a severe and progressive neurologic disease, and symptoms vary depending on disease type. Patients suffer from developmental delays, seizures, skeletal abnormalities, and cognitive decline. Today, treatment options are limited to controlling symptoms.

A recently performed high-throughput screen identified the retinoid compound Tazarotene restoring sulfatase activity in vitro^43^. The investigation of Tazarotene with the KNIME-based ChEMBL Annotation Dashboard (Figure 9) shows that it has so far only been reported active on retinoic acid receptors and the effect on sulfatase activity has not yet been reported in ChEMBL. Tazarotene has a long history being approved since 1997, which suggests a potentially favorable toxicological profile. However, it has only been approved and tested for topical application, and further preclinical and clinical studies are necessary to evaluate its safety and efficacy for systemic administration.

**Figure 9.**
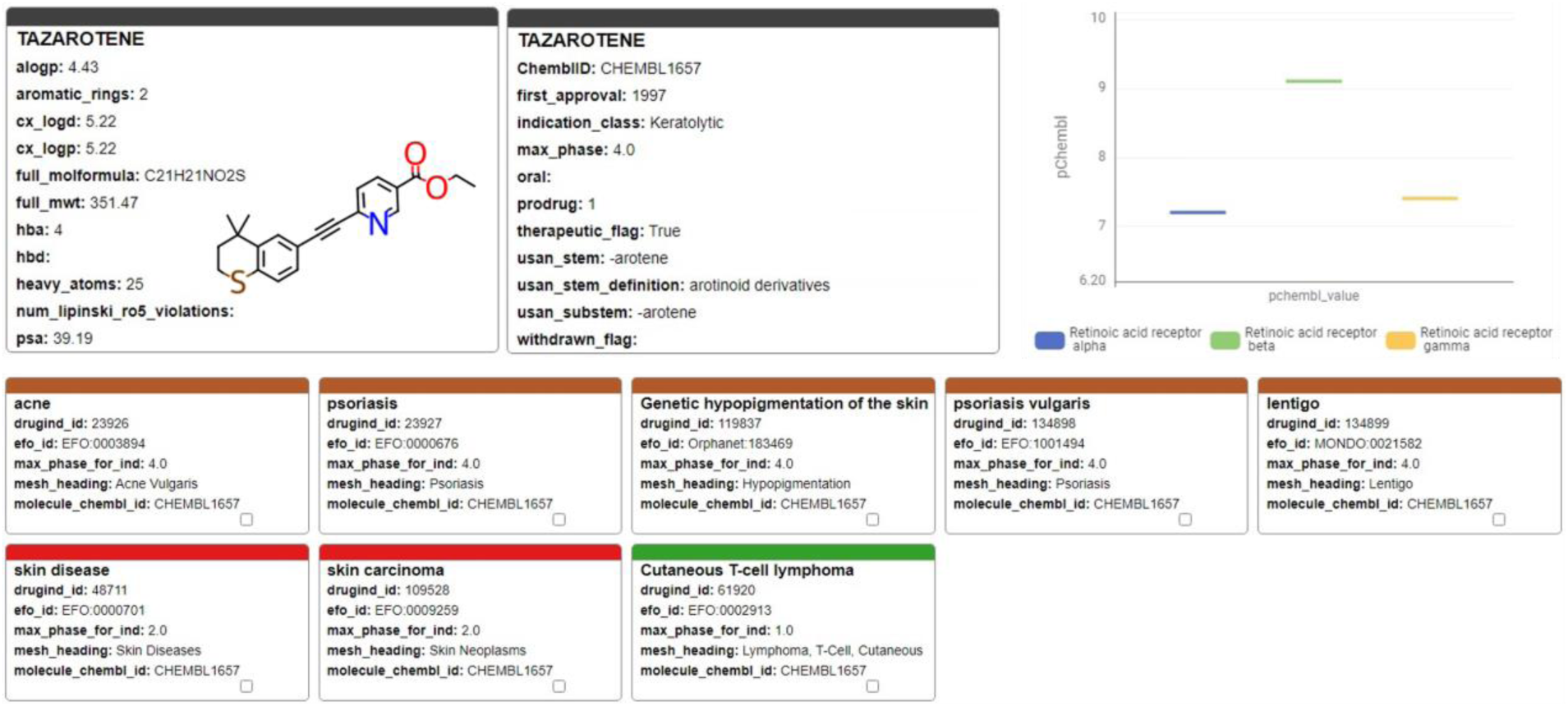
Tazarotene; chemical, pharmaceutical and biochemical information taken from the ChEMBL Annotation Dashboard

#### Use case 2

There are 5 different human Potassium Voltage-gated Channel (Kv7.1 to Kv7.5) with distinct tissue expression patterns and functional duties. These membrane receptors are also known as KCNQ1-5. KCNQ2 is deeply involved in brain development of newborns and responsible for most cases of youth epilepsy events either when its function is inhibited (loss of function) or activated (gain of function). KCNQ2 postnatal genetic screen is commonly run in maternity units. High throughput screening assays run in ITMP using the R4A set, we identified several inhibitors, but also an agonist (JNJ-37822681, CHEMBL3234237) which compete for the same binding site of Retigabine (EZOGABINE, CHEMBL41355) [Carotenuto L. et al. *Repurposing the fast-dissociating D2 antagonist antipsychotic JNJ-37822681 as a neuronal Kv7 channel opener for epilepsy treatment*, submitted]. This MoA identification was only possible through structural and mutations studies defined by annotation-driven hypotheses and structural biology evidence on Retigabine. Furthermore, the structural determinants of JNJ activities prompted us to have deeper insights into the nature of antagonist/agonist profile in KCNQ2 and how selectivity against cardiac KV7 subtypes could be managed, removing critical undesired activities. JNJ-37822681 is not toxic and had already reached Phase II status as D2 antagonist and was withdrawn because it did not reach the clinical potency targets. Annotations of such KCNQ2 modulating compounds (Figure 10), not only the clinical candidate but also other hits found by screening, is of critical importance to assess their clinical potential and to suggest further preclinical studies to eventually bring this molecule to a second indications application. Indeed, for JNJ-37822681, we were able to apply for a patent disclosing its novel second indication [Pat application PCT/EP2024/088161]

**Figure 10.**
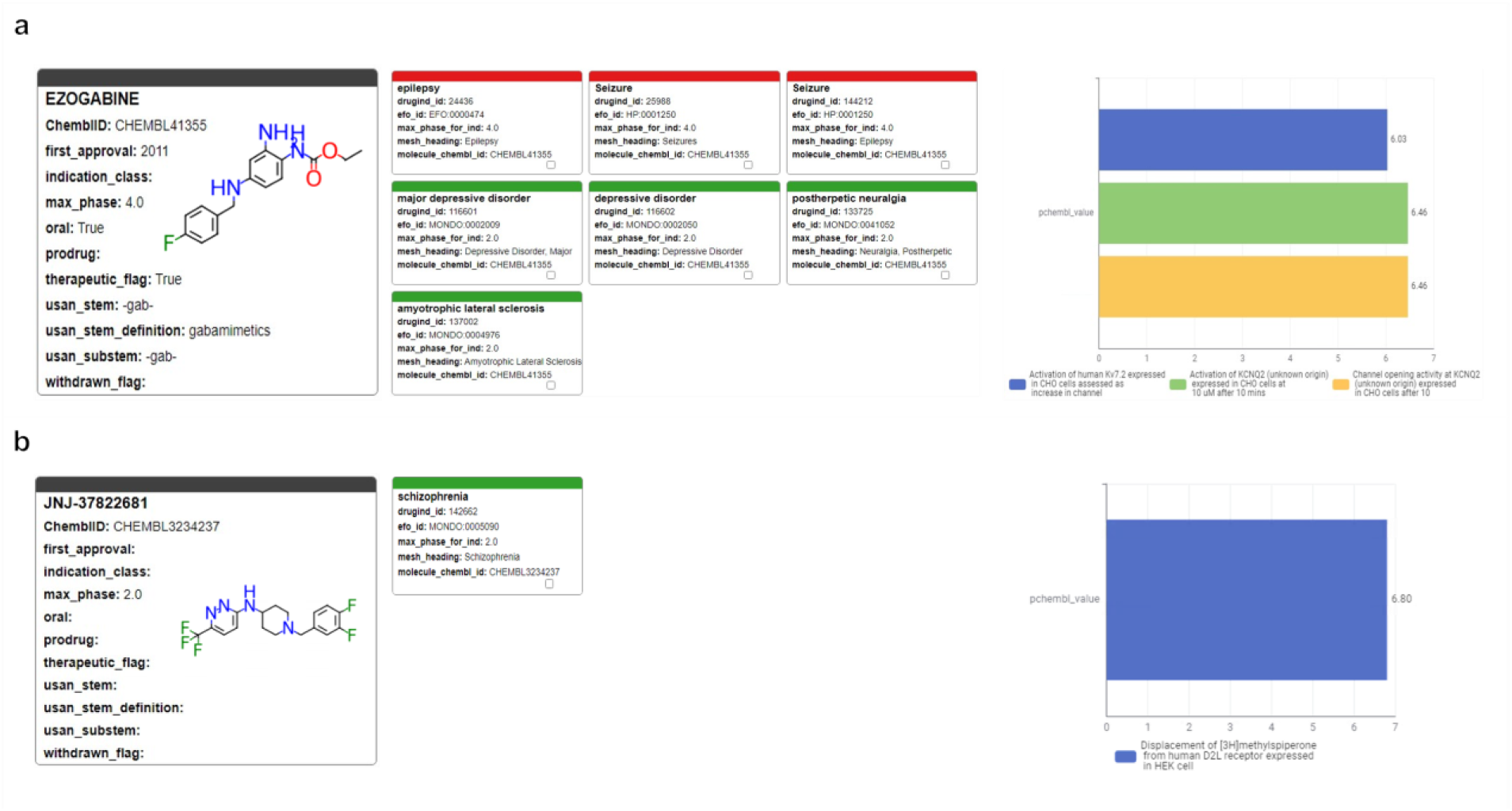
Profiles of pharmaceutical, clinical and biological results retrieved from the ChEMBL Annotation Dashboard for a) Ezogabine (Retigabine) and b) JNJ-37822681.

#### Use case 3

Identifying previously unrecognized mechanisms of action for small molecules is essential for understanding and mitigating adverse effects. In drug discovery, structural similarity plays a pivotal role, particularly in lead optimization, by facilitating the identification of potential compound-target interactions. The Chemical Biology Atlas enables the identification of such interactions by utilizing curated target annotations of structurally related compounds.

To evaluate this approach for target identification, we examined domperidone, a European Medicines Agency (EMA)-approved dopamine D₂ receptor (DRD2) antagonist indicated for gastrointestinal motility disorders (Figure 11). Domperidone is reported to inhibit dopamine D_2_ (DRD2) and dopamine D₃ (DRD3) receptors, and the human Ether-à-go-go-Related Gene (hERG) channel^44^. Pimozide, a structurally related antipsychotic drug (Tanimoto coefficient = 0.58), exhibits a comparable target profile. Notably, pimozide also inhibits the alpha-1 adrenergic receptor (α₁ -AR), with a reported Ki value of 39 nM. Interestingly, a previous computational study predicted α₁ -AR as a target for domperidone and confirmed its binding affinity^45^.

**Figure 11.**
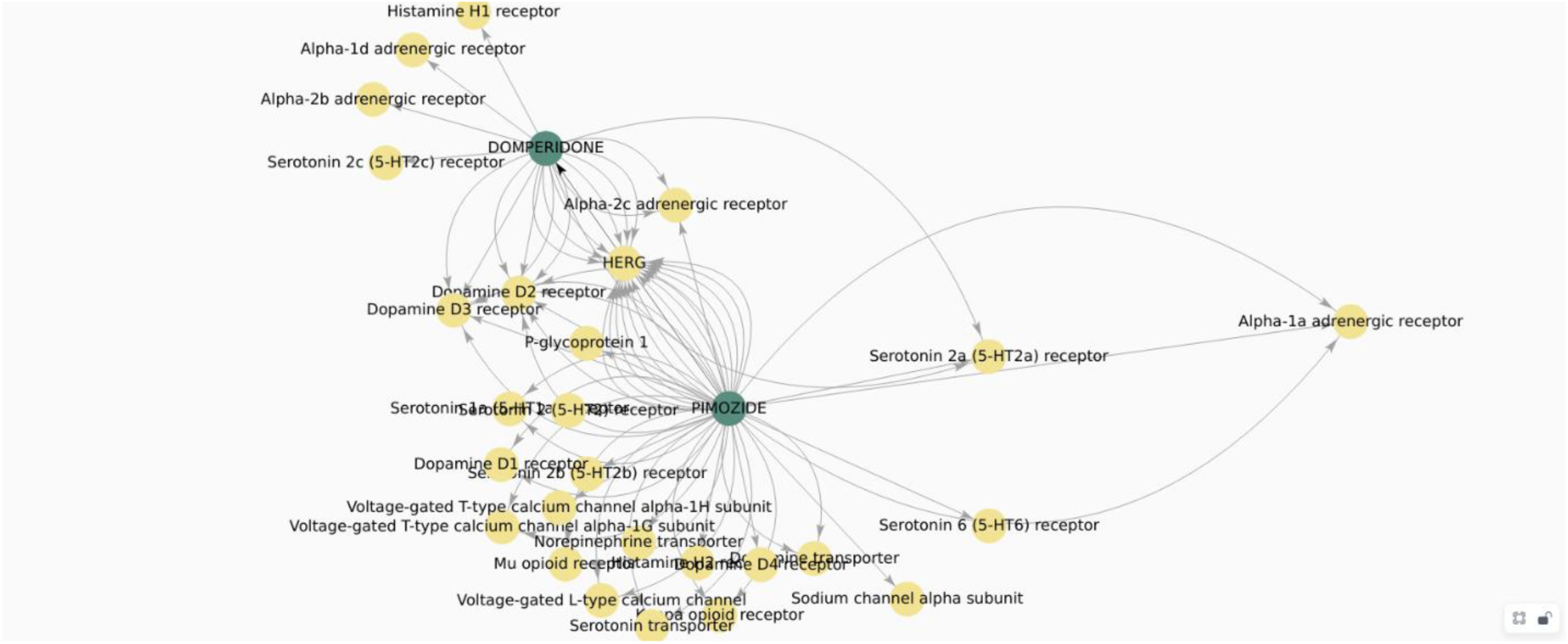
Target space graph plot of domperidone and its analogues generated from the “Compound’s analogue space search” page of the Chemical Biology Atlas.

These findings underscore the utility of our database in identifying potential off-target interactions, thereby enhancing the understanding of adverse effects associated with small molecules.

## Discussion

### Data Analysis and Interpretation of Results

#### Assessing Data Consistency Between KNIME and Python Workflows

A critical component of our analysis involved assessing failure rates for both pipelines. We tracked and inspected data retrieval failures, identifying root causes such as API limitations, connection issues, or database-related problems. A comprehensive analysis of results for the R4A set confirmed that both the KNIME and Python pipelines deliver comparable annotated data, with the Python method generally providing a marginally larger number of results. Importantly, we quantified a strong overlap between the two pipelines’ results, as measured by the Jaccard Index (JI), demonstrating that both the pipelines are robust and reliable (Figures 4-8, Supplementary Tables S1-S2 and Figure S2). Annotation counts for compound-centric data demonstrated close alignment, with Jaccard similarity indexes of 86%, 85% and 80% for data identifiers such as ChEMBL ID, sChEMBL ID and PubChem Compound Identifier (CID), respectively (Figure 4a). This trend continues in the other compound-centric categories, such as compound classification by development phases (Figures 4b-4c) as well as number of compounds with known MoA, indication class, EFO term, and MeSH heading (Figure 4d), where a strong overlap was consistently measured (JI values from 81% to 100 %). Likewise, both methods collected highly similar annotation-centric data, such as the unique number of MoAs, indication classes, MeSH headings, EFO terms (Figure 5a), as well as the number of pChEMBL values, single protein targets and assays (Figure 5b). Similar results were also observed for the distribution of the protein family classification (Figures 6a-b), as well as for the distribution of top-ranked targets with the largest number of associated pChEMBL activities or unique compounds (Figures 7a-b).

#### Interpretation of the annotation results in the context of drug discovery and repurposing

Analysis of the distribution of pChEMBL activities values across different target categories from both pipelines (Figure 6) shows that enzymes dominate the dataset with several measurements exceeding more than twice the number recorded for membrane receptors (Supplementary Table S3). Epigenetic regulators, transcription factors, ion channels and transporters have fewer measurements, while other categories show substantially less activity data. Overall, the distribution of target categories obtained from the two pipelines is highly comparable. Figure 7a shows the top-ranked biological targets based on the number of associated pChEMBL values, highlighting those that have been mostly characterized. Epidermal growth factor receptor, erbB1, and Carbonic anhydrase II emerge as the most heavily studied targets followed by dopamine D2 receptor. Notably, there is a steep decline in measurement counts further down the list, with targets like Carbonic anhydrase I, Serotonin 1a receptor, and various kinases, GPCRs and histone deacetylases in the mid-range (400-800 values). In contrast, Figure 7b ranks targets by the number of unique compounds tested, emphasizing chemical diversity rather than measurement volume. While some overlap exists with Figure 7a, the rankings differ as some targets have many measurements for relatively few compounds while others are associated with a broader spectrum of chemical diversity. This comparison underscores the importance of considering both the quantity of activity data and the diversity of tested compounds when evaluating targets for drug repurposing. The prominence of G protein-coupled receptors (GPCRs) in both rankings is consistent with their well-known pharmacological importance as major drug targets and their central role in drug discovery and repurposing efforts^46,47^.

The top-ranked disease indications or categories by the number of unique compounds per EFO term (Supplementary Figure S2) emphasize the broad therapeutic diversity of the R4A set. There is a particular enrichment for cancer-related pathologies, alongside other major diseases such as cardiovascular, infection and metabolic diseases. To gain insights into the landscape of drug promiscuity within the R4A set, we analyzed the distribution of compounds based on the number of unique targets for which activity is reported in ChEMBL (Figure 8). The observed data shows that compounds interact on average with 5.5 targets, a result which is in line with previous studies of drug promiscuity^48^.

#### How does the R4A set cover the ChEMBL’s druggable proteome?

The identification and characterization of druggable protein targets is central to drug discovery and repurposing. ChEMBL is widely recognized as a comprehensive, curated database of bioactive molecules and their protein targets, and is frequently used as a reference for the druggable proteome due to its extensive coverage and rigorous annotation standards^49,50^. We performed a comparative analysis of the R4A set and ChEMBL to quantitatively assess the overlap of their chemical and druggable spaces (Figures 12-13 and S3-4). We considered the ChEMBL approved drugs and clinical compounds as subset of ChEMBL (v35) space comprising compounds with assigned max_phase (ChEMBL max_phase range between −1 and 4) and relative target data from assay types B and F (binding and functional), with confidence scores ≥ 8 and pChEMBL values ≥ 6.

**Figure 12:**
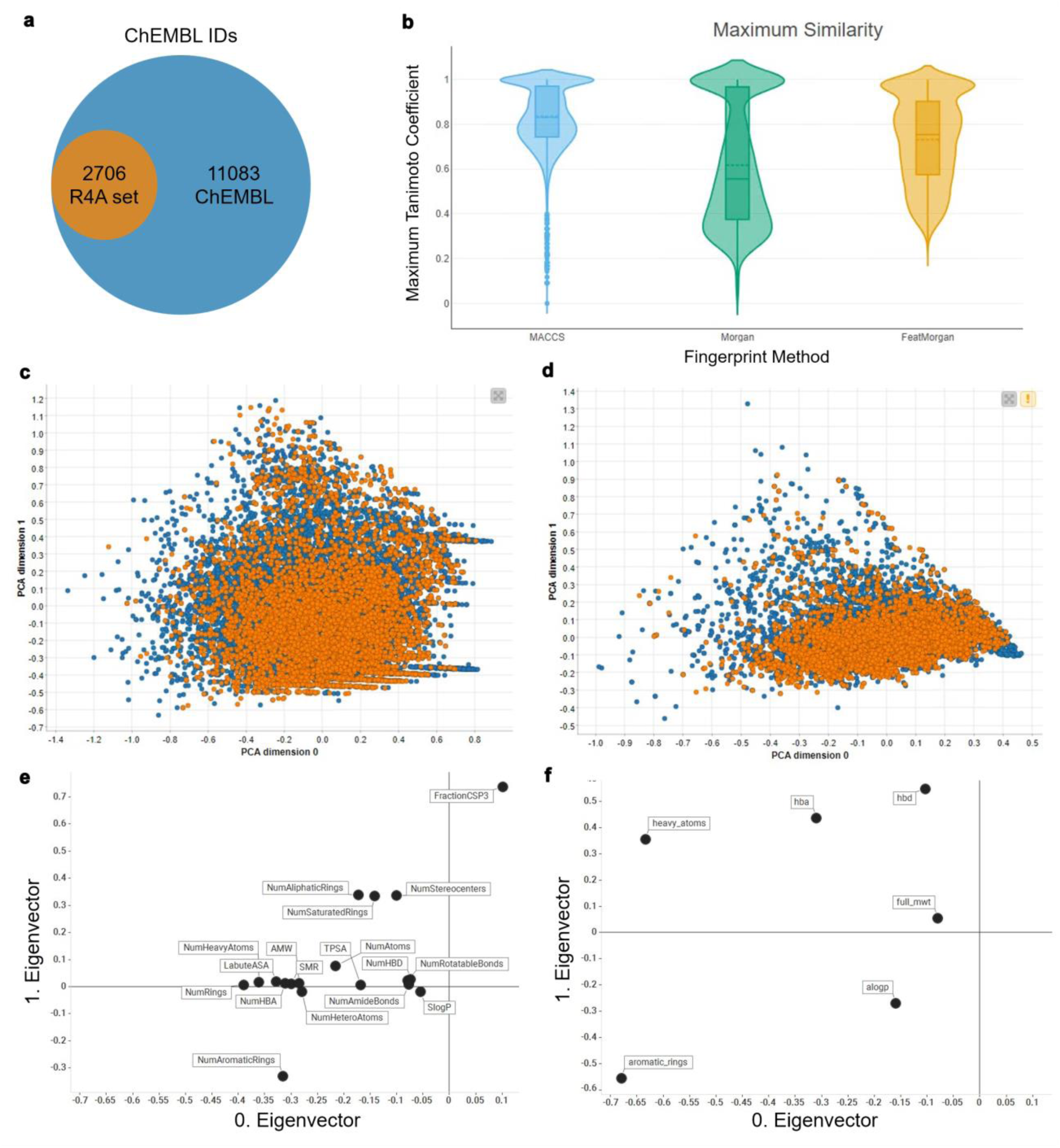
Comparison between the R4A set (orange) and the ChEMBL clinical compounds (blue) compounds; a) R4A set has ∼20 % overlap of ChEMBL IDs with all ChEMBL clinical compounds, b) Violin plots of Maximum Tanimoto similarity of ChEMBL clinical compounds with R4A set using MACCS (blue), Morgan radius 2 (green) and FeatMorgan radius 2 (orange) fingerprints, c) Principal component analysis (PCA) on physico-chemical properties calculated with the RDKit Descriptor Calculation Node in KNIME, d) Physico-chemical properties retrieved from ChEMBL database using the python-based annotation Pipeline. e) Biplot of eigenvectors (loadings) representing magnitude and direction of the feature’s contribution for the PCA using physico-chemical properties calculated with e) the RDKit Descriptor Calculation Node in KNIME, **f**) Physico-chemical properties retrieved from ChEMBL database.

**Figure 13.**
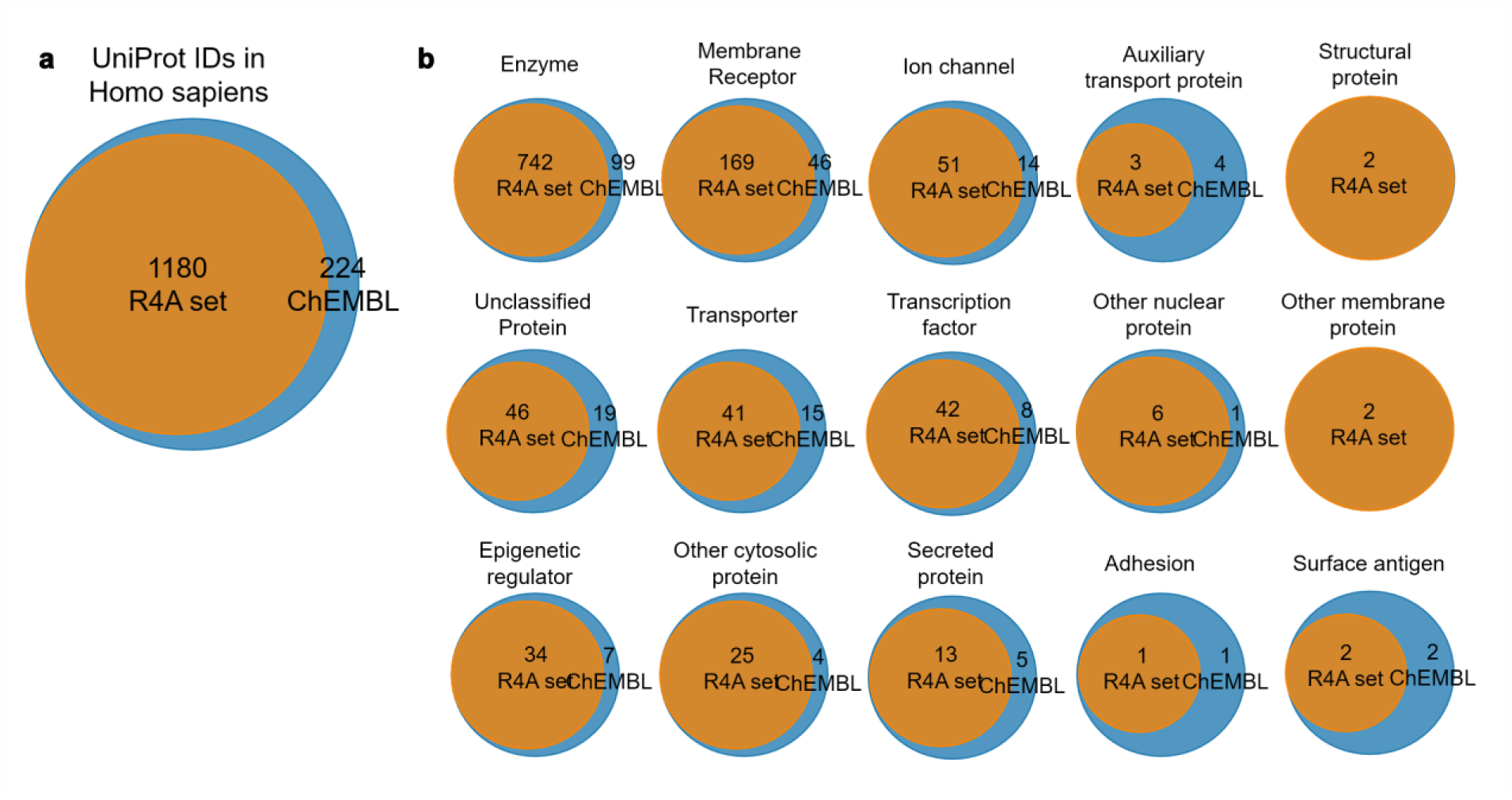
Overlap of druggable proteome in Homo sapiens between the R4A set and ChEMBL chemical compounds for targets associated with clinical compounds. a) Overall overlap of UniProt IDs; b) Overlap UniProt IDs by target class. Count is based on the number of unique UniProt IDs, considering binding and functional assays with a pChEMBL value ≥ 6 and a confidence score ≥ 8. The data shown was obtained from the Python pipeline.

When considering the overlap of ChEMBL IDs, the R4A set constitutes a small subset (∼20%) of the number of the ChEMBL clinical compounds (Figure 12a.However, when comparing both set using molecular fingerprint similarity we identified a substantial degree of structural overlap between compounds of the two sets (Figure 12b). Using MACCS fingerprint similarity, which focusses on substructural features, the majority of ChEMBL clinical compounds appear to be represented in the R4A set. A greater degree of separation is found using Morgan and featMorgan fingerprints, both of which capture circular atom-level environments. The separation is less pronounced using featMorgan fingerprints, which incorporate pharmacophoric properties and therefore emphasizes functional rather than structural similarity.

Employing a principal component analysis (PCA) on the structural properties utilizing common RDKit descriptors revealed a good coverage of the ChEMBL chemical space as shown in Figure 12c. Partitioning of the two dimensional data frame into quadrants of 0.1 x 0.1 gave 304 quadrants occupied for the ChEMBL chemical space, of these 250 are also occupied by the R4A set giving a coverage of 82 % of the ChEMBL chemical space (see Methods section and Supplementary Figure S3. For interpretation of the contribution of the original features, the PCA loadings, we visualize the eigenvectors of the two principal components in Figure 12e, showing the greatest magnitude for number of sp3 hybridization. The details of the chemical space analysis can be found in the methods section. Similarly, and shown in Figure 12d when using the physico-chemical descriptors retrieved (alogp, number of aromatic rings, molecular weight, hydrogen bond acceptors and donors and number of heavy atoms) from the ChEMBL database using the annotation pipelines a good correlation can be seen, with a coverage of 78% (from 158 quadrants). Here, the biplot of the eigenvectors of the components reveals the number of aromatic ring system as dominant feature. The stripes evident in the PCA plot in Figure 12e are most likely due to the abundance of integer descriptors in the dataset.

A broad coverage of the target landscape was measured considering the overlap of UniProt IDs in all organisms (∼77%, Supplementary Figure S4a) and Homo Sapiens in particular (∼84%, Figure 13a). A substantial overlap was also measured when considering the protein targets grouped by target class (Supplementary Figure S4b and Figure 13b). However, results highlighted the potential to extend the R4A set by systematically incorporating molecules with a mechanism of action towards targets that are currently not well represented or even absent.

#### How does the R4A set cover the Drug Repurposing Hub?

The Broad Institutes’ Drug Repurposing Hub is a curated database covering pharmaceutical and biochemical information such as MoA, target proteins, disease indication and is a rich source of information in the field of drug repurposing. The R4A set is designed as a subset of the Broad institutes’ collection. We investigated the Drug information table (version 3/24/2020) downloadable from the Drug Repurposing Hub covering structural information on >20,000 samples (not unique active chemicals, e.g. different salt forms, vendors, batch IDs). A comparison of the two collections based on molecular fingerprints, e.g. MACCS, Morgan and FeatMorgan shows a general close similarity (Figure 14a). A principal component analysis utilising the RDKit descriptors as described above revealed that our institutes’ R4A set covers 95 % of the chemical space of the Drug Repurposing Hub collection (Figures 14b and S5). This underscored the width of diversity the R4A set covers and the richness of information our pipelines produced for investigating the potential of drug repurposing candidates also outside the R4A set.

**Figure 14.**
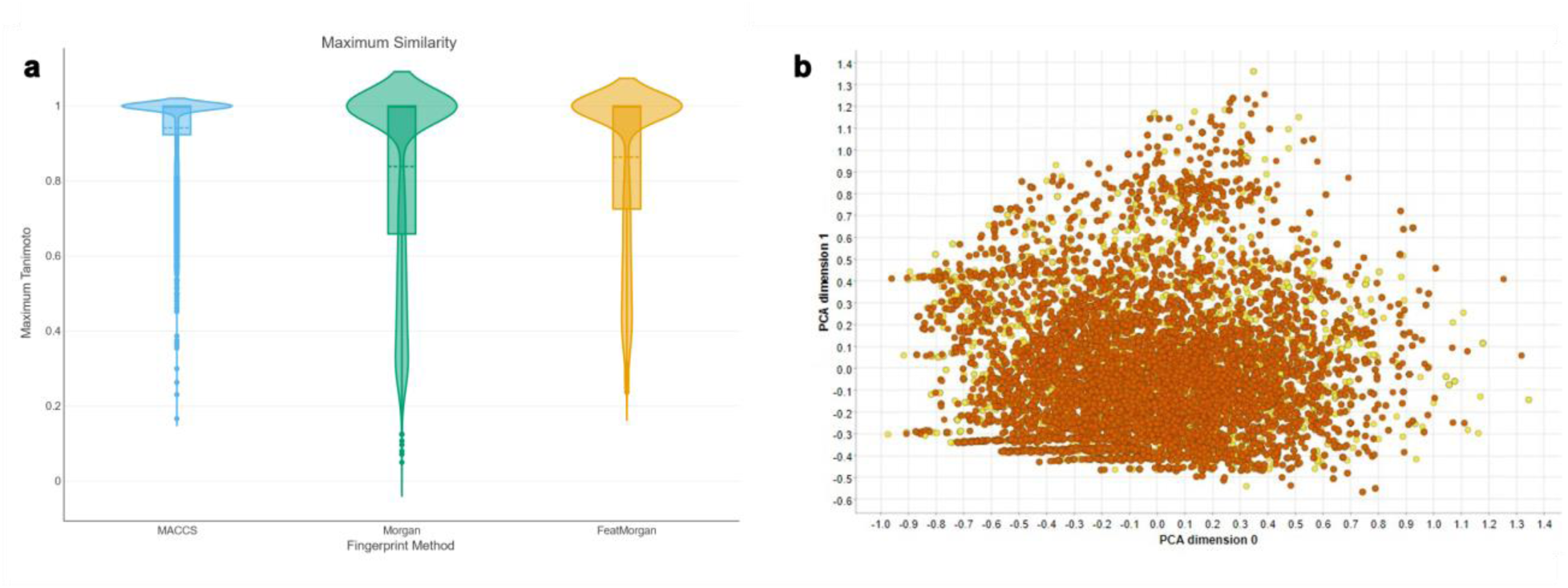
Comparison between the R4A set (orange) and Drug Repurposing Hub (yellow). a) Violin plots of Maximum Tanimoto similarity of Drug Repurposing Hub compounds with R4A set using MACCS (blue), Morgan radius 2 (green) and FeatMorgan radius 2 (orange) fingerprints, b) Principal component analysis (PCA) on physico-chemical properties calculated with the RDKit Descriptor Calculation Node in KNIME.

### Practical Implications

The integration of KNIME- and Python-based annotation pipelines, complemented by interactive dashboards, offers a robust and callable solution for streamlining data-driven drug repurposing research. By automating the extraction and integration of chemical and bioactivity data from heterogeneous public sources, these tools facilitate a task that is generally labor-intensive and error prone.

Validated on a harmonized subset of the Specs repurposing collection (over 5000 compounds), the pipelines demonstrated reproducibility and adaptability, making them suitable for any user-provided compound library, ensuring data quality and traceability in all types of data driven drug discovery and repurposing studies.

The interactive dashboards—KNIME’s ChEMBL Annotation Dashboard and the Neo4j-powered Chemical Biology Atlas— allow multilayered exploration of chemical, pharmacological, and relational data.

At the time of writing, the R4A Annotation Tool was available on KNIME HUB for 4 months, reaching ∼ 80 downloads, while the latest Dashboard version was online for 1 month reaching > 20 downloads. This demonstrates a clear interest of the scientific community in these tools.

Together, we think that these tools can streamline informed decision-making in drug repurposing, facilitating translational research by bridging computational and experimental data.

### Future implementations

The comparative evaluation of KNIME and Python workflows revealed complementary strengths as well as directions for future development. One key improvement will be the expansion of input flexibility, allowing users to query the pipelines using additional identifiers such as drug names or CAS numbers, e.g. Further integration with additional databases will enrich the depth of annotation and provide a more comprehensive biological context. Here, the Remedi4All in-silico drug repurposing catalogue^51^ will serve as guideline. Analytical capabilities can also be expanded, particularly within the KNIME dashboard, to include exploratory data analysis techniques such as clustering, PCA, UMAP, t-SNE, scaffold extraction and descriptor-based profiling to support deeper analyses into chemical space. The integration of commercial vendor database for the identification of structurally related analogues could be implemented to support structure-activity relationship (SAR) analyses, aiding the progression of an initial hit compound to a more advanced lead substance.

While the two workflows are not intended to guide clinical decision-making, they provide a framework for exploration of bioactive compounds and hypothesis generation in drug repurposing.

In conclusion, this study presents a harmonized and reproducible framework for automated annotation and analysis of compound libraries, particularly for drug repurposing initiatives. The KNIME and Python pipelines enable systematic integration of curated chemical and bioactivity data from public repositories, in alignment with FAIR principles. The accompanying dashboards facilitate intuitive exploration of complex datasets, supporting both compound-centric and target-centric analyses. The framework is adaptable to any user-provided compound library and is accessible to researchers with diverse computational backgrounds. Although the current focus is on compounds with clinical trial history, the methodology is broadly applicable to various drug discovery contexts.

## Methods

### Ingestion of data from public resources

For our annotation workflow, we leveraged several open-source biological databases and their programmatic access tools. PubChem^52^, maintained by the NIH, houses over 111 million unique chemical structures and 271 million bioactivity data points derived from 1.2 million biological assays^53^. We leveraged PubChem’s identifier exchange service to ground and map chemicals using a controlled vocabulary, ensuring consistency in chemical representation. Additionally, we integrated data from ChEMBL^54^, which contains detailed information on 2.5 million drug-like compounds and their biological effects^55^. Due to its semi-automated data aggregation methods, ChEMBL serves as a pivotal resource in our annotation workflow, providing reliable and comprehensive drug discovery-related data. By programmatically accessing these databases, our workflow efficiently taps into rich repositories of biological and medicinal chemistry information, ensuring robust and accurate annotations.

### Programmatic language used by the workflows

Our primary objective with the workflows is to provide technical solutions for the dynamic annotation of collections with relevant, up-to-date bioactivity and clinical results. The goal is to ensure that these annotations are easily reusable by scientists with varying levels of informatics expertise. To achieve this, we designed our annotation workflows using two programmatic languages: KNIME and Python, catering to a broader bioinformatics and cheminformatics community. Konstanz Information Miner KNIME^40^ is an open-source, low-code platform for creating visual workflows tailored to data mining, analysis, machine learning, and data visualization. KNIME offers a vast library of pre-built “nodes” for data transformation, cleaning, statistical analysis, visualization, deployment, and retrieval. Additionally, it supports numerous extensions for cheminformatics, bioinformatics, REST services, and various scripting languages such as Python, R, Java, and SQL. The KNIME Community Hub serves as a rich repository of freely available workflows, featuring more than 27,000 community-shared workflows at the time of writing.

The KNIME workflows were created with KNIME Analytics Platform version 5.4.2. and uses the following extensions: KNIME Base nodes, KNIME Excel Support, KNIME Expressions, KNIME Javasnippet, KNIME JSON-Processing, KNIME Math Expression (JEP), KNIME REST Client Extension, KNIME XML-Processing, KNIME Base Chemistry Types and Nodes, KNIME JavaScript Views, KNIME Quick Forms, KNIME SVG Support, KNIME Views, KNIME-CDK, KNIME Plotly and RDKit Nodes Feature.

Python^56^ is a widely used, high-level programming language known for its simplicity, versatility, and extensive ecosystem of scientific libraries. In this project Python was used to develop the Chemical Annotator pipeline. The pipeline primarily integrates the *chembl_resource_client*^57^ and the *pubchempy*^58^ libraries, in combination with in-house functions that directly query RESTful endpoints to access ChEMBL, PubChem and KEGG databases. To support data processing and integration, the workflow also employs several well-established libraries, including *pandas* for data manipulation, *requests* for API communication, *math* and *json* for data formatting and transformation.

### Minting of identifier

When acquiring a compound library, each organization assigns internal identifiers for reference and internal use. However, when compound libraries are shared across organizations, these internal identifiers lose relevance. To ensure consistency and enable FAIR data principles, standardized identifiers are essential. To address this need for consistent identifiers across organizations, we developed a KNIME workflow that assigns unique identifiers to compounds within the shared collection.

The identifier generation process begins with compound cleaning using the RDKit node in KNIME. This cleaning step selectively retains counter ions that are physiologically relevant in aqueous test solutions (Figure 15). For instance, counter ions such as sodium, chlorine, and acetate are removed, while lithium and antimony are retained. Unique identifiers are assigned based on InChIKeys, leveraging their detailed and unique representation of chemical structures. All enantiomers of the same chemical scaffold are considered unique entities; however, they share a common prefix in the generated identifier, allowing easy identification without requiring a similarity search. Additionally, mixtures and drug combinations are assigned both a unique combination identifier and individual identifiers for each component. The complete KNIME workflow for identifier generation is available on KNIME Hub^59^.

**Figure 15.**
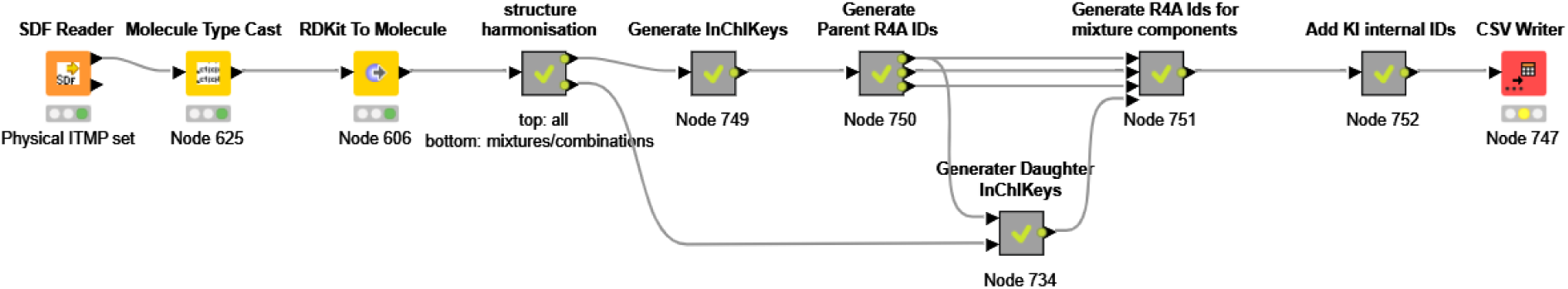
Layout of the structure for identifier minting workflow published on KNIME Community Hub.

**Figure 16.**
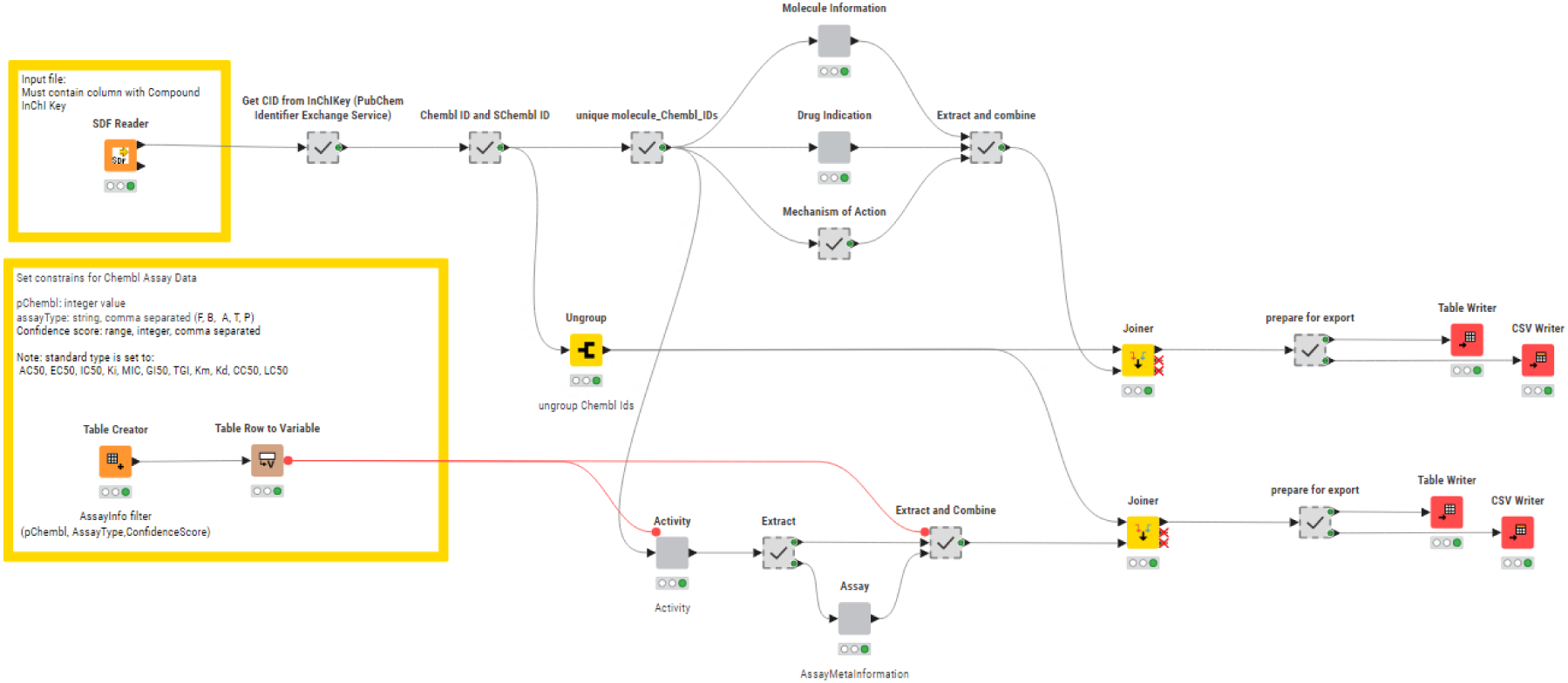
Layout of the structure for KNIME annotation workflow published on KNIME Community Hub^60^.

### KNIME Annotation Workflow

The InChiKey representation generated from the structure harmonisation pipeline are used as input in the annotation workflow. These are used in a “GET Request” node use as input for retrieving PubChem Compound Identifier (CID) from the PubChem Identifier Exchange service. The CIDs are then used as input for a second request to the PubChem Identifier Exchange service for retrieving compound synonyms. During the development we observed inconsistencies in output quantity which was caused by server timeouts leading to empty response bodies. We were able to minimise this problem by introduction of recursive loops on empty response occurrences. The Synonyms list for each CID includes, besides others, ChEMBL IDs and sChEMBL IDs, of which the ChEMBL IDs are used for querying the ChEMBL Database in five separate metanodes. The first metanode queries the *molecule* table retrieving chemical and physicochemical properties e.g. molecular weight, number of aromatic rings, molecular formula, max phase, indication class, first year of approval. The second metanode queries the *drug_indication* table and retrieves clinical trail information such as efo terms, mesh headings, indications and the maximum clinical trial phase reached. The third metanode queries the *mechanism* table with information about the mechanism of action of the molecule. All information gathered from these three metanodes is combined and forms the MedChem output. The fourth metanode queries the *activity* table retrieving assay results such as pChEMBL values, target, assay description and assay ID. The assay ID is used in the fifth metanode for querying the *assay* table extracting information on cell type or assay organism and assay confidence score. The information from these final two metanodes is combined to form the BioAssay output. To improve performance and speed, requests are bundled for 20 compounds in case of the activity table and 200 compounds in other requests and up to 100 loop recursions in case of empty response bodies. The ChEMBL REST API default limit for responses is 20 and is set to the maximum 200 in all GET requests. The total number of entries is extracted from each response and is used for calculating the number of iterations with increased offset numbers.

### Python Annotation Pipeline

The Chemical Annotator is composed of five scripts — *chemical_annotator.py*, *chembl_utils.py*, *kegg.py*, *misc_utils.py* and *pubchem_utils.py* (Supplementary Figure S1).

The *chemical_annotator.py* script serves as command-line interface and execution driver for the entire pipeline. It begins by parsing arguments for input and output file paths and the type of chemical identifier to use (SMILES, InChI, or InChIKey). It sets up a logging system to track the execution process and any errors. Once the input CSV file is read into a DataFrame, the script calls the “process_compounds” function from *misc_utils.py* which acts as a high-level orchestrator for compound-level data aggregation. By calling appropriate functions defined in the *chembl_utils.py*, *pubchem_utils.py* scripts, it ensures that each compound is enriched with as much relevant chemical and biological information as possible. The function handles missing data and provides a real-time progress bar, making it suitable for large-scale processing.

The *chembl_utils.py* contains a suite of functions that interact with the ChEMBL web services^61^. For compound-level data, a series of functions convert chemical identifiers into ChEMBL IDs, retrieve compound metadata, therapeutic uses and biological mechanisms. For assay-level data, functions filter assay information and associated publication metadata by user-defined confidence score and pchembl value. For target-level data, functions retrieve target metadata, and protein classification. These functions are designed to be modular and fault-tolerant. The results are saved into Excel files. After this, the script proceeds to collect assay and target data associated with the compounds that have been found in ChEMBL.

EC numbers are then mapped to KEGG pathways (function defined in *kegg.py* script). The final output includes reports on drug information, assay results, mechanism of action, target data and pathway annotations.

In the present study, the SMILES representation generated from the structure harmonisation pipeline has been considered as input.

### Neo4J Database and NeoDash

A Neo4j graph database was developed with a schema (Figure 17) reflecting the drug-related knowledge collected by the Chemical Annotator. At the core of the graph, are *Compound* nodes, each annotated with detailed chemical and metadata properties such as compound_id, smiles, canonical_smiles, first approval, max_phase, and molecule_synonym among others. Compound nodes are connected to *Target* nodes which include attributes like target_chembl_id, UNiProt_ID, target_pref_name, target_organism and protein_hierarchy, capturing both biological entity and classification. The graph also includes nodes for *Indication_class, Mesh_Heading*, and *EFO_Term*, each representing therapeutic categories, biomedical indexing terms, and ontology-based disease descriptors, respectively.

**Figure 17.**
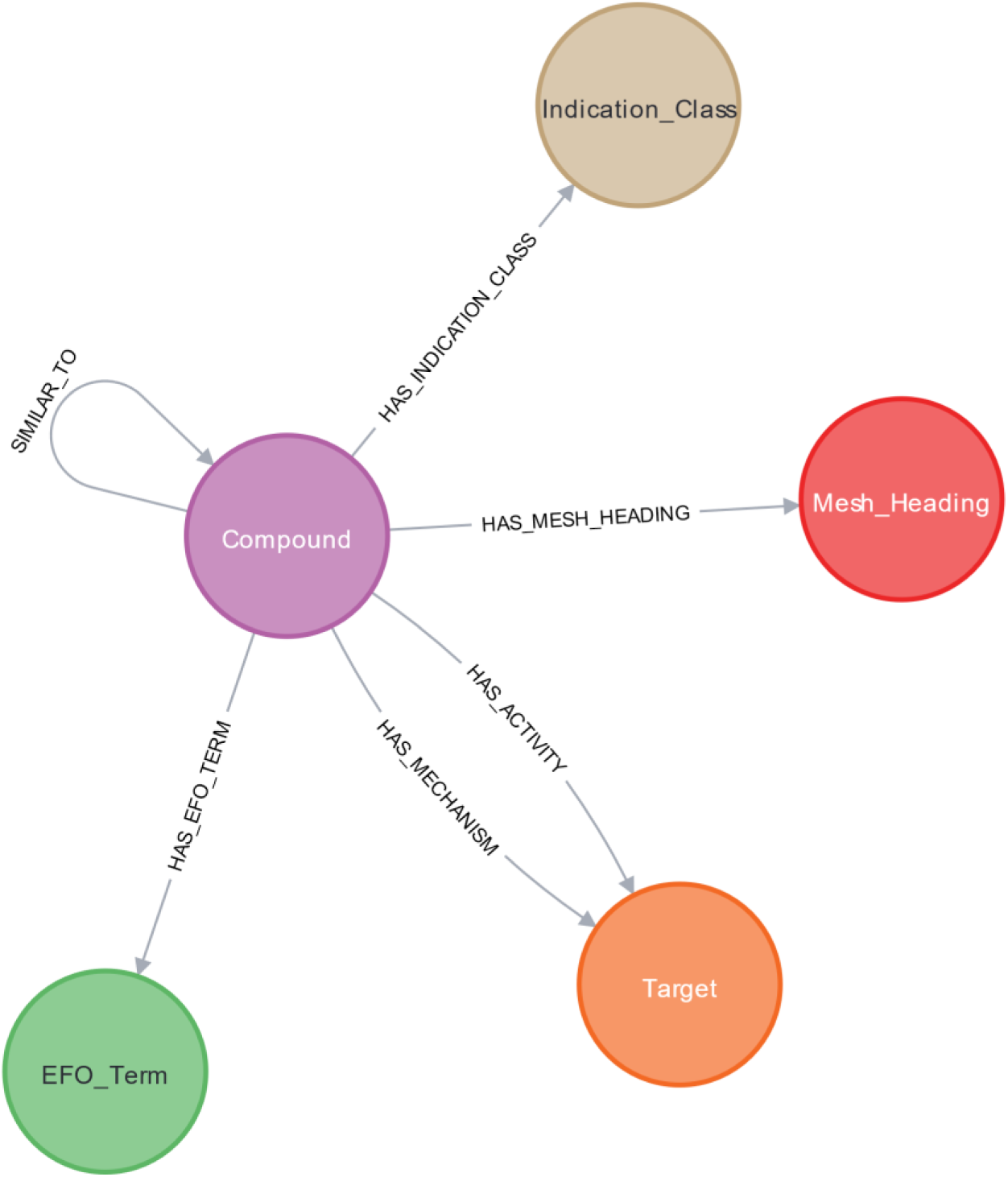
Neo4j schema.

Relationships between these nodes are richly annotated: HAS_ACTIVITY links *compound* nodes to *target* nodes with assay metadata such as assay_type, standard_type, confidence_score, pchembl_value, and doi, HAS_MECHANISM captures mechanistic insights through action_type and mechanism_of_action, SIMILAR_TO quantifies compound similarity using the Tanimoto coefficient (Tc).

Other relationships like HAS_EFO_TERM, HAS_MESH_HEADING and HAS_INDICATION_CLASS provide links to disease and classification ontologies. This integrated graph structure serves as an engine behind an interactive dashboard built using NeoDash, enabling intuitive exploration, querying, and visualization of drug-related knowledge.

### Fingerprint similarity

For comparison of the chemical space of the R4A library and the ChEMBL clinical compounds, input SMILES were used to calculate molecular fingerprints of types MACCS, Morgan and featMorgan using RDKit fingerprint node. The circular fingerprints, Morgan and feature-based Morgan, were generated using a radius of 2 and a bit vector length of 1024. The Tanimoto coefficients are calculated using the Fingerprint Similarity Node of the CDK KNIME extension. Here, for each of the ChEMBL clinical set compounds the maximum Tanimoto similarity to the closest R4A set analogue is retrieved.

Likewise, all pairwise similarities implemented in the Neo4j database — amounting to over 13 million— were precalculated using RDKit with Morgan fingerprints (radius 2, 1024-bit vectors), based on the SMILES representations of the R4A set.

### Principal component analyses

The principal component analyses were performed using KNIME analytics platform. Input components are either physico-chemical properties retrieved by the KNIME annotation workflow or calculated from canonicalized SMILES using the “RDKit Descriptor Calculation” node. All properties were normalized to 0 – 1 scale using the “Normalizer” node before entering the “PCA” node, and the dataset is reduced to two dimensions. The two-dimensional data frame is sliced into quadrants of 0.1 x 0.1 for dimension 0 and dimension 1 and quadrants occupying either of the reference library (ChEMBL clinical set or Drug Repurposing Hub) are considered. The proportion of considered quadrants that are also occupied by at least one compound of the R4A set is determined. To perform this calculation, datapoints for each dimension are rounded to one decimal place, a “GroupBy” node is used to group the data set by rounded dimension 0 and dimension 1 and number of datapoints for each compound collection for each group is counted. The chemical space coverage is calculated as percentage of quadrants occupied also for the R4A set of the total chemical space covered by the reference library (Supplementary Figures S3 and S5). For determination of the PCA loadings, the “PCA Compute” node is used to calculate the eigenvalues of the first two components (second output port).

## Data Availability

Demonstrator data used in this study can be accessed on Zenodo at https://doi.org/10.5281/zenodo.16359229.

## Code Availability

The KNIME workflow and dashboard are available on KNIME HUB.

| KNIME Workflow | Link |
| --- | --- |
| Annotation Dashboard | https://hub.knime.com/fraunhoferitmp/spaces/Public/Remedi4All/Chembl%20Annotation%20Dashboard~GG1R6X_poNP7r852/current-state |
| Annotation Workflow | https://hub.knime.com/fraunhoferitmp/spaces/Public/Remedi4All/R4A_Annotation_Tool_v3.1~em5BEui1IMPof_Ry/current-state |
| Identifier generation | https://hub.knime.com/fraunhoferitmp/spaces/Public/Remedi4All/R4A_StructureHarmonisation_IDgenerator~9qa2Cmp7a47KU5GR/current-state |

The python code of the Chemical Annotator workflow is available on GitHub and Zenodo. The Chemical Biology Atlas is accessible on the SciLifeLab Serve platform.

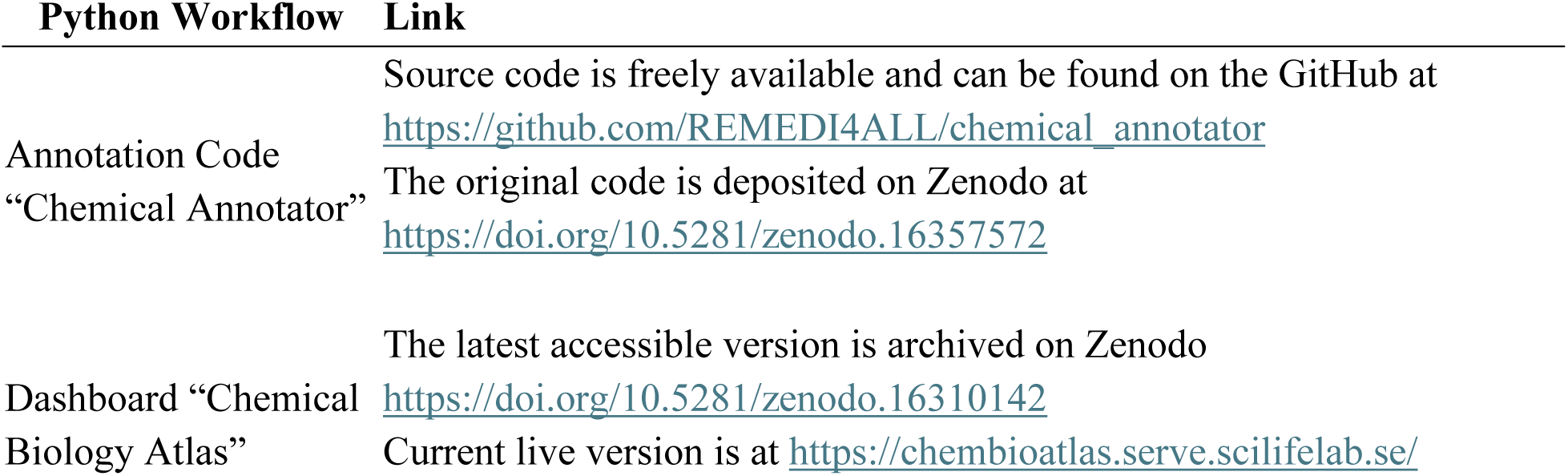

## Author contributions

J.R.: Conceptualization, Methodology, Development of the KNIME annotation pipeline and the KNIME dashboard, Data Analysis, Writing-Reviewing and Editing, Visualization; F.B.: Conceptualization, Supervision, Methodology, Development of the Python annotation pipeline, Data Analysis, Development of the NEO4J database and NeoDash dashboard, Writing-Reviewing and Editing; B.S.L.: Data analysis, Visualization, Writing-Reviewing and Editing. Y.G.: Writing-Reviewing and Editing; A.L.G.: Writing-Reviewing and Editing; Z.T.: Writing-Reviewing and Editing; T.A.: Writing-Reviewing and Editing; J.H.: Writing-Reviewing and Editing; A.J.J.: Writing-Reviewing and Editing; P.G.: Writing-Reviewing and Editing; A.Z.: Writing-Reviewing and Editing.

## Acknowledgements

We would like to thank the authors of the public resources used in our work for making their datasets available to the scientific community. The REMEDi4ALL project has received funding from the European Union’s Horizon Europe Research & Innovation programme under grant agreement No 101057442. The Chemical Biology Consortium Sweden (CBCS), node KI, is a national research infrastructure funded by the Swedish Research Council (dr.nr.2021-00179) and SciLifeLab. Technical infrastructure for hosting the Chemical Biology Atlas was provided by SciLifeLab Serve (https://serve.scilifelab.se), a platform developed and supported by SciLifeLab Data Centre. The work was also funded by the Research Council of Finland (No. 351507 to ZT). This work reflects only the authors’ view, and the EC is not responsible for any use that may be made of the information it contains.

## Notes

### Competing Interest Statement

The authors have declared no competing interest.

## References

1. Pushpakom, S. et al. Drug repurposing: progress, challenges and recommendations. Nat. Rev. Drug Discov. 18, 41–58 (2019).

2. Pinzi, L., Bisi, N. & Rastelli, G. How drug repurposing can advance drug discovery: challenges and opportunities. Front. Drug Discov. 4, 1460100 (2024).

3. Cha, Y. et al. Drug repurposing from the perspective of pharmaceutical companies. Br. J. Pharmacol. 175, 168–180 (2018).

4. Chow, W. A., Jiang, C. & Guan, M. Anti-HIV drugs for cancer therapeutics: back to the future? Lancet Oncol. 10, 61–71 (2009).

5. Teo, S. K. et al. Thalidomide in the treatment of leprosy. Microbes Infect. 4, 1193–1202 (2002).

6. Palumbo, A. et al. Thalidomide for treatment of multiple myeloma: 10 years later. Blood 111, 3968–3977 (2008).

7. Guilbert, S. M., Cardoso, D., Lévy, N., Muchir, A. & Nissan, X. Hutchinson-Gilford progeria syndrome: Rejuvenating old drugs to fight accelerated ageing. Methods 190, 3–12 (2021).

8. Jarada, T. N., Rokne, J. G. & Alhajj, R. A review of computational drug repositioning: strategies, approaches, opportunities, challenges, and directions. J. Cheminformatics 12, 46 (2020).

9. Oprea, T. I. & Overington, J. P. Computational and Practical Aspects of Drug Repositioning. ASSAY Drug Dev. Technol. 13, 299–306 (2015).

10. Lee, S. et al. High-throughput identification of repurposable neuroactive drugs with potent anti-glioblastoma activity. Nat. Med. 30, 3196–3208 (2024).

11. Tanoli, Z. et al. Computational drug repurposing: approaches, evaluation of in silico resources and case studies. Nat. Rev. Drug Discov. (2025) doi:10.1038/s41573-025-01164-x.

12. Corsello, S. M. et al. The Drug Repurposing Hub: a next-generation drug library and information resource. Nat. Med. 23, 405–408 (2017).

13. Gaulton, A. et al. ChEMBL: a large-scale bioactivity database for drug discovery. Nucleic Acids Res. 40, D1100–D1107 (2012).

14. Wang, Y. et al. PubChem: a public information system for analyzing bioactivities of small molecules. Nucleic Acids Res. 37, W623–W633 (2009).

15. Wishart, D. S. et al. DrugBank: a knowledgebase for drugs, drug actions and drug targets. Nucleic Acids Res. 36, D901–D906 (2008).

16. Skuta, C. et al. Probes & Drugs portal: an interactive, open data resource for chemical biology. Nat. Methods 14, 759–760 (2017).

17. Harding, S. D. et al. The IUPHAR/BPS Guide to PHARMACOLOGY in 2024. Nucleic Acids Res. 52, D1438–D1449 (2024).

18. Wang, L. et al. The landscape of the methodology in drug repurposing using human genomic data: a systematic review. Brief. Bioinform. 25, bbad527 (2024).

19. Jin, Q., Leaman, R. & Lu, Z. PubMed and beyond: biomedical literature search in the age of artificial intelligence. eBioMedicine 100, 104988 (2024).

20. De Rosa, M. C., Purohit, R. & García-Sosa, A. T. Drug repurposing: a nexus of innovation, science, and potential. Sci. Rep. 13, 17887, s41598-023-44264–7 (2023).

21. Otero-Carrasco, B. et al. Identifying patterns to uncover the importance of biological pathways on known drug repurposing scenarios. BMC Genomics 25, 43 (2024).

22. Masoudi-Sobhanzadeh, Y., Omidi, Y., Amanlou, M. & Masoudi-Nejad, A. DrugR+: A comprehensive relational database for drug repurposing, combination therapy, and replacement therapy. Comput. Biol. Med. 109, 254–262 (2019).

23. Drug repurposing: approaches, methods and considerations | Elsevier. www.elsevier.com https://www.elsevier.com/industry/drug-repurposing.

24. Bang, D., Lim, S., Lee, S. & Kim, S. Biomedical knowledge graph learning for drug repurposing by extending guilt-by-association to multiple layers. Nat. Commun. 14, 3570 (2023).

25. Wilkinson, M. D. et al. The FAIR Guiding Principles for scientific data management and stewardship. Sci. Data 3, 160018 (2016).

26. Gadiya, Y. et al. FAIR data management: what does it mean for drug discovery? Front. Drug Discov. 3, 1226727 (2023).

27. Fortier, I. et al. Maelstrom Research guidelines for rigorous retrospective data harmonization. Int. J. Epidemiol. dyw075 (2016) doi:10.1093/ije/dyw075.

28. Weiner, M. W. et al. Impact of the Alzheimer’s Disease Neuroimaging Initiative, 2004 to 2014. Alzheimers Dement. 11, 865–884 (2015).

29. Beam, A. L. & Kohane, I. S. Big Data and Machine Learning in Health Care. JAMA 319, 1317 (2018).

30. Lehne, M., Sass, J., Essenwanger, A., Schepers, J. & Thun, S. Why digital medicine depends on interoperability. Npj Digit. Med. 2, 79 (2019).

31. Index - FHIR v5.0.0. https://www.hl7.org/fhir/.

32. Heinzke, A. L. et al. A compound-target pairs dataset: differences between drugs, clinical candidates and other bioactive compounds. Sci. Data 11, 1160 (2024).

33. Huang, Y. et al. DrugRepoBank: a comprehensive database and discovery platform for accelerating drug repositioning. Database 2024, baae051 (2024).

34. Gan, J. et al. DrugRep: an automatic virtual screening server for drug repurposing. Acta Pharmacol. Sin. 44, 888–896 (2023).

35. Narwani, T. J., Srinivasan, N. & Chakraborti, S. NOD: a web server to predict New use of Old Drugs to facilitate drug repurposing. Sci. Rep. 11, 13540 (2021).

36. Srinivasan, B. & Lloyd, M. D. Quantitation and Error Measurements in Dose–Response Curves. J. Med. Chem. 68, 2052–2056 (2025).

37. SPECS-factsheet-repurposing library.pdf.

38. Corsello, S. M. et al. The Drug Repurposing Hub: a next-generation drug library and information resource. Nat. Med. 23, 405–408 (2017).

39. Neo4j Graph Database & Analytics – The Leader in Graph Databases. Graph Database & Analytics https://neo4j.com/ (2025).

40. Berthold, M. R. et al. KNIME: The Konstanz Information Miner. in Data Analysis, Machine Learning and Applications (eds. Preisach, C., Burkhardt, H., Schmidt-Thieme, L. & Decker, R.) 319–326 (Springer Berlin Heidelberg, Berlin, Heidelberg, 2008). doi:10.1007/978-3-540-78246-9_38.

41. Remedi4All Homepage. REMEDi4ALL https://remedi4all.org/.

42. Kanehisa, M. KEGG: Kyoto Encyclopedia of Genes and Genomes. Nucleic Acids Res. 28, 27–30 (2000).

43. Schlotawa, L., et al. Drug screening identifies tazarotene and bexarotene as therapeutic agents in multiple sulfatase deficiency. EMBO Mol. Med. 15, e14837 (2023).

44. Barone, J. A. Domperidone: A Peripherally Acting Dopamine_2_ -Receptor Antagonist. Ann. Pharmacother. 33, 429–440 (1999).

45. Keiser, M. J. et al. Predicting new molecular targets for known drugs. Nature 462, 175–181 (2009).

46. Hauser, A. S., Attwood, M. M., Rask-Andersen, M., Schiöth, H. B. & Gloriam, D. E. Trends in GPCR drug discovery: new agents, targets and indications. Nat. Rev. Drug Discov. 16, 829–842 (2017).

47. Ballante, F., Kooistra, A. J., Kampen, S., De Graaf, C. & Carlsson, J. Structure-Based Virtual Screening for Ligands of G Protein–Coupled Receptors: What Can Molecular Docking Do for You? Pharmacol. Rev. 73, 1698–1736 (2021).

48. Hu, Y., Gupta-Ostermann, D. & Bajorath, J. EXPLORING COMPOUND PROMISCUITY PATTERNS AND MULTI-TARGET ACTIVITY SPACES. Comput. Struct. Biotechnol. J. 9, e201401003 (2014).

49. Heinzke, A. L. et al. A compound-target pairs dataset: differences between drugs, clinical candidates and other bioactive compounds. Sci. Data 11, 1160 (2024).

50. Zdrazil, B. et al. The ChEMBL Database in 2023: a drug discovery platform spanning multiple bioactivity data types and time periods. Nucleic Acids Res. 52, D1180–D1192 (2024).

51. idrc-r4a.com. https://www.idrc-r4a.com/landing.

52. PubChem. PubChem. https://pubchem.ncbi.nlm.nih.gov/.

53. Kim, S. et al. PubChem in 2021: new data content and improved web interfaces. Nucleic Acids Res. 49, D1388–D1395 (2021).

54. ChEMBL - ChEMBL. https://www.ebi.ac.uk/.

55. Zdrazil, B. et al. The ChEMBL Database in 2023: a drug discovery platform spanning multiple bioactivity data types and time periods. Nucleic Acids Res. 52, D1180–D1192 (2024).

56. Welcome to Python.org. Python.org https://www.python.org/ (2025).

57. chembl/chembl_webresource_client. Chemical Biology Services @ EMBL-EBI (2025).

58. Swain, M. mcs07/PubChemPy. (2025).

59. R4A_StructureHarmonisation_IDgenerator – fraunhoferitmp. KNIME Community Hub https://hub.knime.com/fraunhoferitmp/spaces/Public/Remedi4All/R4A_StructureHarmonisation_IDgenerator~9qa2Cmp7a47KU5GR/current-state (2023).

60. R4A_Annotation_Tool_v3.1 – fraunhoferitmp. KNIME Community Hub https://hub.knime.com/fraunhoferitmp/spaces/Public/Remedi4All/R4A_Annotation_Tool_v3.1~em5BEui1IMPof_Ry/current-state (2025).

61. Davies, M. et al. ChEMBL web services: streamlining access to drug discovery data and utilities. Nucleic Acids Res. 43, W612–W620 (2015).

