## Supplementary Information for "From Library to Landscape: Integrative Annotation Workflows for Compound Libraries in Drug Repurposing"

Number of pages: 8
Number of tables: 3
Number of figures: 5

### Table of contents

Tables S3-S4

Figures S4-S8

**Table S1.** Compound-centric information retrieved by the KNIME and Python annotation pipelines**.**

| **Description** | **KNIME annotation** | **Python annotation** | **Overlap** | **Jaccard index[a]** |
|---|---|---|---|---|
| Total library size | 5254 | 5254 | 5254 | 1 |
| Unique compounds after structure harmonisation | 5230 | 5230 | 5230 | 1 |
| CID | 5091 | 5217 | 4592 | 0.8 |
| sChEMBL ID | 4585 | 4787 | 4296 | 0.85 |
| ChEMBL ID | 4688 | 4954 | 4457 | 0.86 |
| Launched | 1155 | 1304 | 1113 | 0.83 |
| Phase 3 | 292 | 302 | 278 | 0.88 |
| Phase 2 | 790 | 812 | 772 | 0.93 |
| Phase 1 | 131 | 133 | 126 | 0.91 |
| Early Phase 1 | 3 | 3 | 3 | 1 |
| Unknown phase | 148 | 150 | 141 | 0.9 |
| Preclinical/discovery | 2169 | 2523 | 2098 | 0.81 |
| Withdrawn | 81 | 90 | 79 | 0.86 |
| clinical trials (total) | 1216 | 1250 | 1179 | 0.92 |
| Compounds with known MoA | 1209 | 1280 | 1156 | 0.87 |
| Compounds with known indication class | 983 | 1114 | 948 | 0.83 |
| Compounds with EFO term | 1851 | 2013 | 1786 | 0.86 |
| Compounds with Mesh heading | 1851 | 2013 | 1786 | 0.86 |

[a] Jaccard Index = $\frac{Ni}{(Na+Nb-Ni)}$

**Table S2.** Annotation-centric information retrieved by the KNIME and Python annotation pipelines. Constrains: pChembl ≥ 6, Confidence score ≥ 8

| Description | KNIME Annotation | Python Annotation | Overlap | Jaccard Index [a] |
|---|---|---|---|---|
| Number of MoA | 630 | 647 | 619 | 0.94 |
| Number of indication class | 348 | 389 | 342 | 0.87 |
| Number of EFO term | 1760 | 1870 | 1739 | 0.92 |
| Number of Mesh heading | 1507 | 1599 | 1489 | 0.92 |
| Endpoints (pChEMBL) | 59751 | 60597 | 57923 | 0.93 |
| Targets | 2202 | 2295 | 2172 | 0.93 |
| Assays | 33913 | 36492 | 33042 | 0.88 |

[a] Jaccard Index $= \frac{\boldsymbol{Ni}}{(\boldsymbol{Na}+\boldsymbol{Nb}-\boldsymbol{Ni})}$

**Table S3.** Target hierarchy counts from the KNIME and Python workflows. Table is available as supplementary file TableS3.xlsx

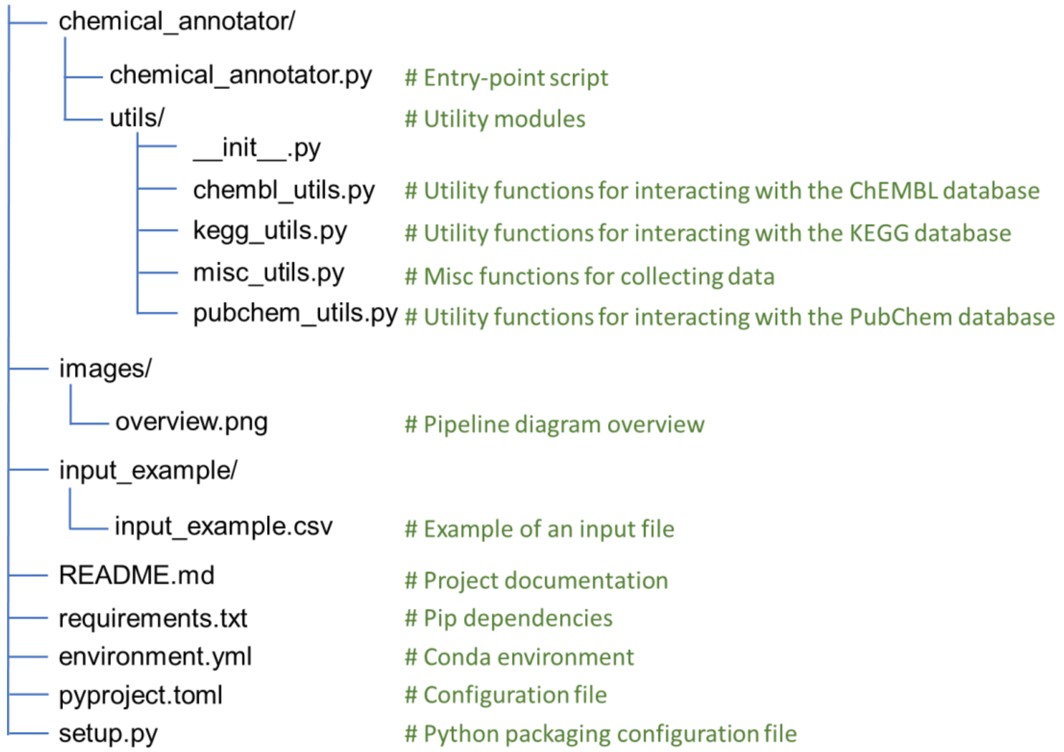


**Figure S1.** Directory structure of the Python annotation pipeline. Comments on the right provide a brief explanation of each file.

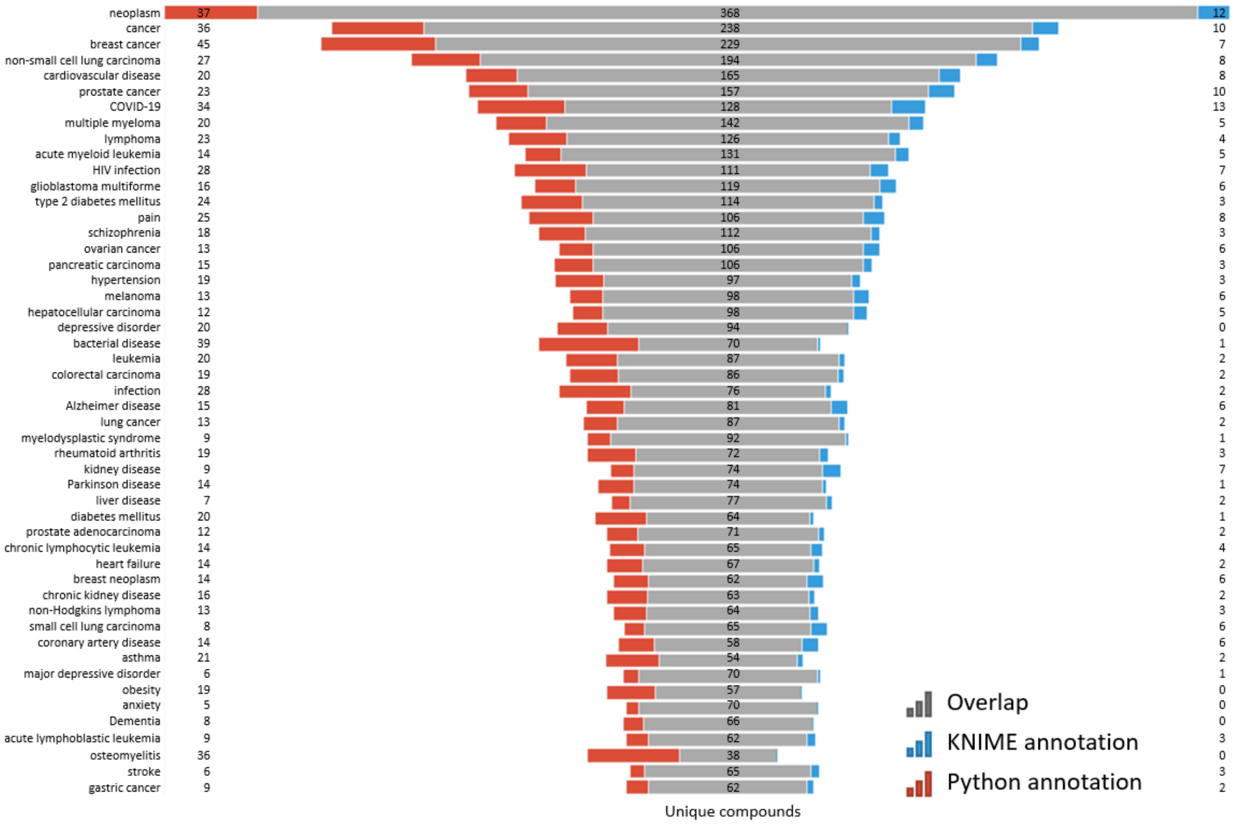


**Figure S2.** Top-ranked EFO terms based on n of unique compounds. Data from binding and functional assays with a pChEMBL value ≥ 6 and a confidence score ≥ 8 were considered.

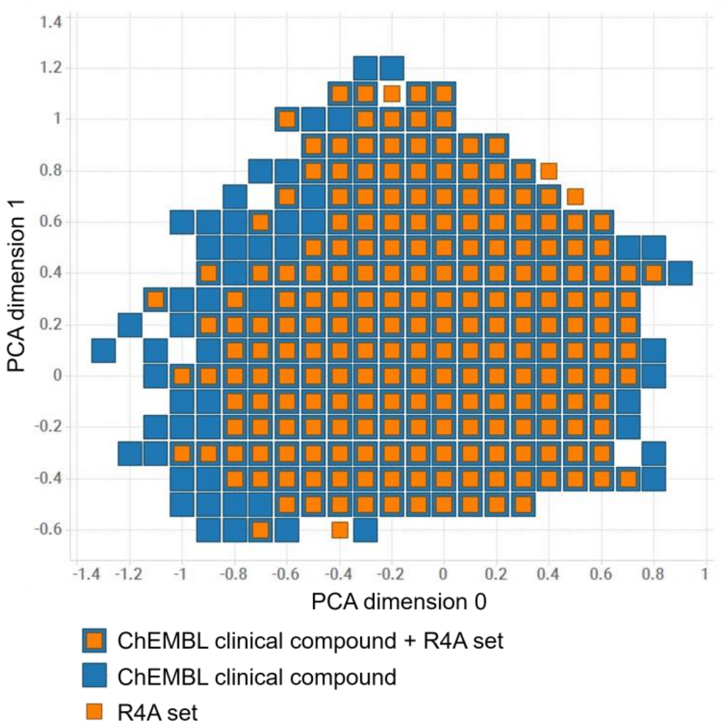


**Figure S3.** Chemical space coverage of R4A set (orange) compared to ChEMBL clinical compounds (blue). Differences in sizes of squares are for better visualization only and do not reflect sector size or compound count.

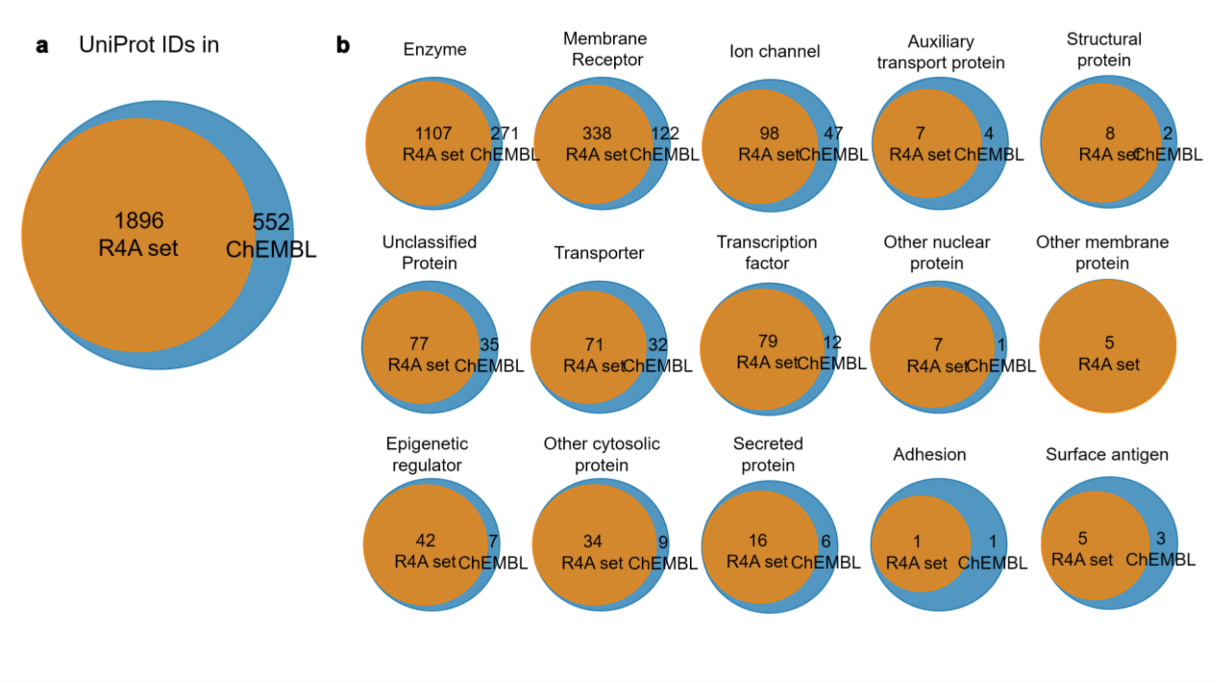


**Figure S4.** Overlap of protein targets between the R4A set and ChEMBL, grouped by target class. a) Overlap of UniProt IDs for targets associated with the molecules (ChEMBL IDs); b) Overlap of UniProt IDs in Homo sapiens. Count is based on the number of unique UniProt IDs, considering binding and functional assays with a pChEMBL value ≥ 6 and a confidence score ≥ 8. The data shown was obtained from the Python pipeline.

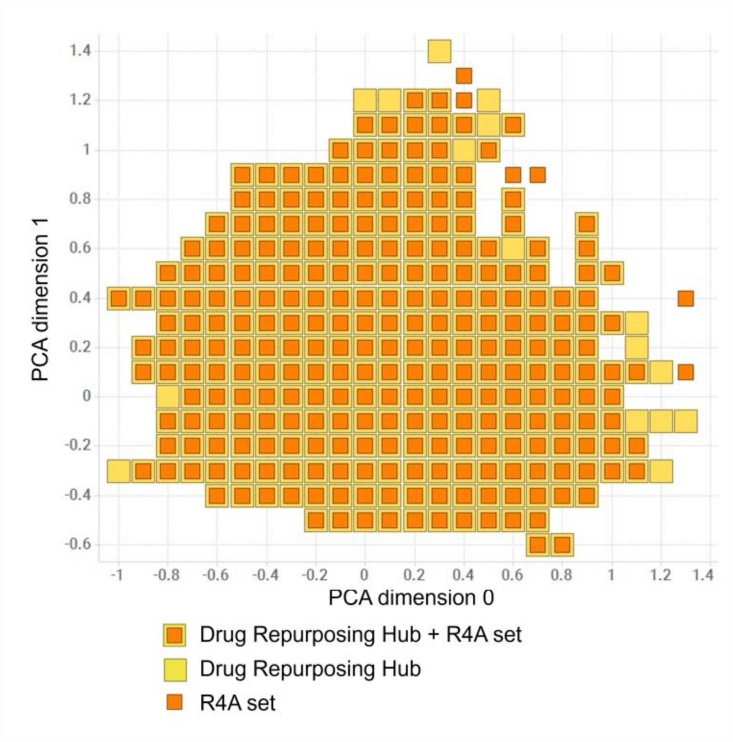


**Figure S5.** Chemical space coverage of R4A set (orange) compared to Drug repurposing hub (yellow). Differences in sizes of squares are for better visualization only and do not reflect sector size or compound count.
